# Single-cell analysis identifies concurrent type 2 and interferon programs in experimental eosinophilic esophagitis

**DOI:** 10.64898/2026.11.18.767761

**Authors:** Anish Dsilva, Kristina Handler, Tali Bar-On, Michal Itan, Shmulik Avlas, Israel Zan-Bar, Chen Varol, Marc E. Rothenberg, Isabelle C. Arnold, Ariel Munitz

## Abstract

**Background & Aims:** Eosinophilic esophagitis (EoE) is a chronic, type 2 cytokine-driven allergic disease of the esophagus. How the immune and structural cell landscape of the esophagus changes between healthy and inflamed tissue requires further investigation.

**Methods:** CD45-enriched esophageal cells from naive and oxazolone (OXA)-challenged wild-type mice were profiled by single-cell RNA sequencing. A cytokine-gated gene expression score identified type 2 program-high cells. Findings were compared with published human esophageal datasets, validated histologically, and examined in *Il13ra1*^- /-^ mice.

**Results:** A total of 62,245 esophageal cells were analyzed. ILC2s accounted for most type 2 cytokine-expressing lymphocytes in naive mice, whereas Th2 cells accounted for approximately 80% following OXA challenge. ILC2s concurrently lost Il5, Il13 and Gata3 expression while retaining Areg. Resident memory-like CD8⁺ T cells became the largest lymphoid population. Mast cells switched from a connective tissue-type to a mucosal-type and partly cycling state, which was confirmed by increased MCPT1⁺ cells. Esophageal eosinophils displayed a signature resembling skin eosinophils of atopic dermatitis rather than gastrointestinal eosinophils. *Il13ra1^-/-^* mice were protected from epithelial proliferation, thickening and eosinophilia. An interferon-associated program characterized lymphoid, myeloid and structural compartments, and its epithelial component was conserved in human EoE.

**Conclusions:** Th2 cells and mucosal-type mast cells are the dominant sources of type 2 cytokines in experimental EoE. This study further identifies a conserved interferon-associated program that accompanies the type 2 response in the immune and structural compartments of the esophagus and highlights it as a candidate contributor to EoE pathology.

**Synopsis:** Single-cell profiling of experimental eosinophilic esophagitis identifies Th2 cells and mucosal mast cells as the dominant sources of type 2 cytokines and reveals a concurrent interferon-associated program that is conserved in human disease.

## Introduction

EoE is a chronic, food antigen-induced T helper type 2 (Th2)-associated, allergic- and immune-mediated disease. It is characterized histologically by eosinophil-predominant mucosal inflammation and clinically by tissue remodeling and esophageal dysfunction^1, 2^. Molecular studies identified a disease-associated transcriptional signature that includes the eosinophil chemoattractant CCL26, linking tissue inflammation to altered epithelial gene expression^3, 4^. The importance of type 2 cytokines (e.g., IL-4 and IL-13) is further supported by the clinical efficacy of IL-4 receptor α (IL-4Rα) blockade with dupilumab, which improves histologic disease and dysphagia^5, 6^. Nonetheless, eosinophil depletion with benralizumab produced a marked histologic response without a corresponding improvement in dysphagia compared with placebo^7^. This dissociation highlights the need to define the cellular responses accompanying eosinophilia and esophageal tissue abnormalities that persist beyond eosinophil accumulation.

Single-cell RNA sequencing (scRNA-seq) has expanded our understanding of the cellular heterogeneity of human EoE. Studies of blood and esophageal T cells identified disease-associated regulatory T (Treg) cells and effector T helper 2 (Th2) cells expressing IL-5 and IL-13^8, 9^. Separate analyses revealed heterogeneous mast cell populations with proliferative and inflammatory features^10^, as well as inflammatory eosinophils^11^. More recently, an integrated human esophageal atlas identified changes in immune and structural cell states and predicted interactions involving fibroblasts, Th2 cells, and mast cells^12^. These observations establish that EoE involves several interacting cellular compartments^13^. Despite these findings, several constraints are intrinsic to what human esophageal biopsies can provide. For instance, control tissue is obtained either from individuals undergoing endoscopy for other indications, who are likely, not disease-free. Thus, neither provides an unambiguous baseline against which disease-associated cellular states can be measured. Furthermore, human biopsies also sample the superficial mucosa of a patchy disease, recover limited numbers of cells per fragment, and capture the lamina propria inconsistently, which constrains analysis of the fibroblast and endothelial compartments in which remodeling occurs. Finally, patients are sampled at a single clinical moment that reflects an unknown disease duration and prior therapy. Experimental models of EoE are useful to overcome these limitations and complement human studies^14^. Experimental models addresses each of these constraints, since naive animals provide a true baseline, the entire esophagus including its structural compartments can be profiled, the timing and dose of antigen exposure are defined, and the genotype can be manipulated so that cellular composition and its cytokine dependence are measured in the same system^15^.

We have recently developed an experimental model of EoE that closely resembles human disease including the infiltration of intraepithelial eosinophils, epithelial remodeling and fibrosis^16^. Furthermore, transcriptome analysis revealed marked overlap between human disease and experimental EoE. Like human studies, experimental EoE was dependent on IL-13Rα1 and TSLP but not on IL-5 and eosinophils^16, 17^. Herein, we aimed to define the cellular and transcriptional landscape at the terminal endpoint of experimental EoE. Esophageal cells from naive mice and challenged mice were profiled by scRNA-seq, and selected findings were examined histologically. Our analyses identified that experimental EoE is characterized by a Th2-associated cytokine expression program, a prominent mucosal mast-cell response. We further identify increased representation of resident memory-like CD8^+^ T cells that is associated with an interferon-signature that is conserved in human patients. These data suggest that the type 2 response in EoE is accompanied with a conserved interferon-associated program, which may contribute to the esophageal pathology in EoE and may persist despite type 2-directed therapy.

## Results

### Single-cell profiling resolves immune and structural cell populations in experimental EoE

To define the cellular landscape of experimental eosinophilic esophagitis, we performed single-cell RNA sequencing of CD45-enriched esophageal cell suspensions from naive and oxazolone (OXA)-challenged wild-type mice. OXA-challenged esophagi were collected on day 36, at the terminal experimental endpoint (Fig. 1A). Following quality control and predicted doublet removal, 62,245 cells were retained, comprising 33,544 cells from naive mice and 28,701 cells from OXA-challenged mice (Fig. 1B).

**Figure 1.**
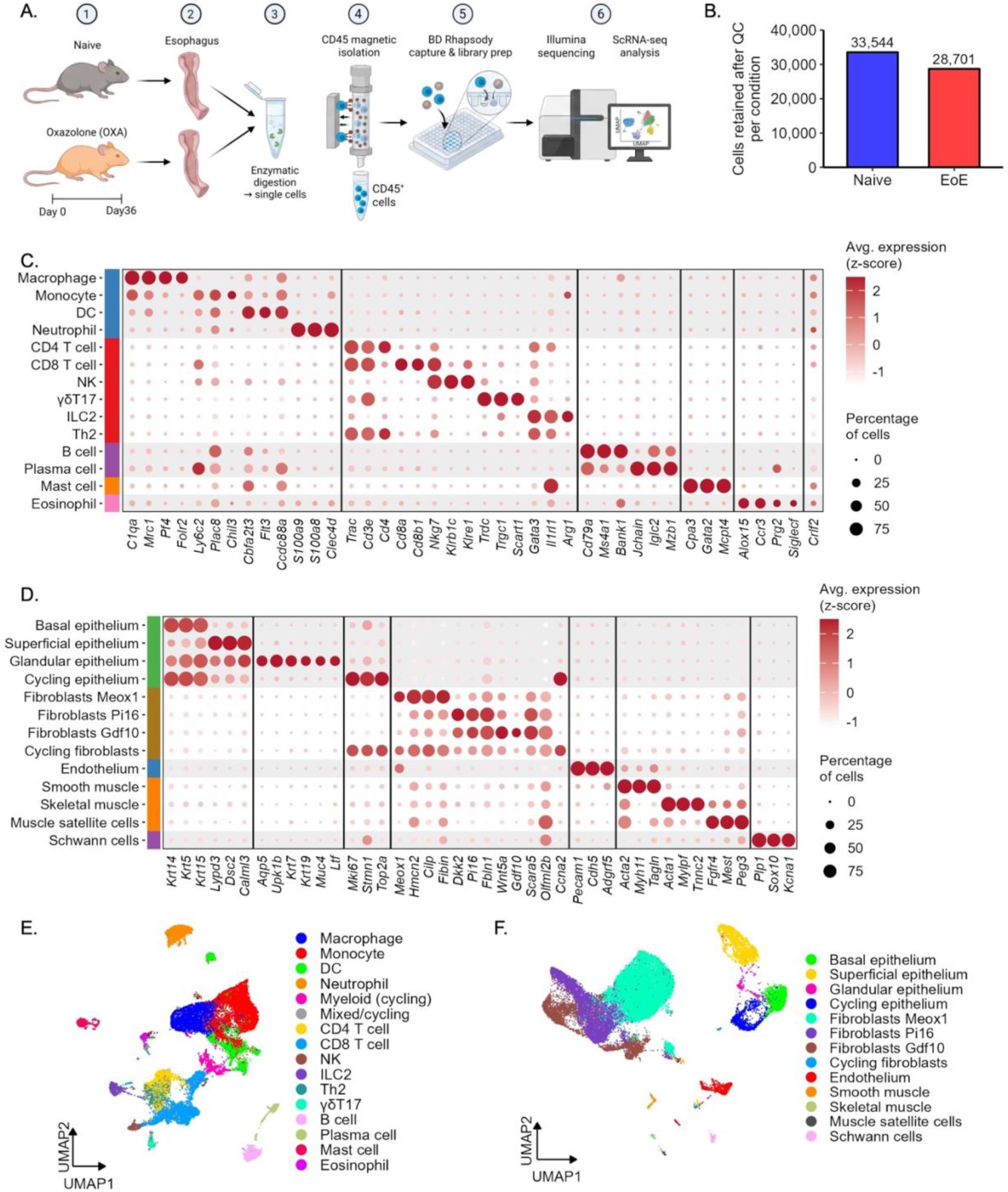
Single-cell RNA sequencing defines the immune and structural cell landscape of experimental EoE. Schematic representation of the experimental design (A). Experimental EoE was induced in wild type (WT) mice using oxazolone (OXA). On day 36, esophagi were obtained from naive and OXA-challenged mice, pooled (2-4 mice per sample; two samples per condition), enzymatically digested, and CD45⁺ cells were enriched and subjected to single-cell RNA sequencing. The number of cells retained after quality control and doublet removal is shown (B). Expression of marker genes in immune (C) and structural (D) cell types is shown. Dot color represents the average expression (z-score) and dot size the percentage of cells expressing each gene (C-D). UMAP representations of immune (E) and structural (F) cells are presented.

Clustering and marker-based annotation identified 34,946 immune cells and 26,928 structural cells. An additional 371 cells with mixed transcriptional profiles were excluded from downstream analyses. The immune compartment comprised myeloid and lymphoid populations, including macrophages, monocytes, dendritic cells, neutrophils, eosinophils, mast cells, T cells, natural killer cells, ILC2s, B cells, and plasma cells (Fig. 1C, 1E). Despite CD45 enrichment, non-immune cells were also recovered, enabling analysis of structural cells such as epithelial cells and fibroblasts (Fig. 1D, 1F). Together, these data establish a cellular and transcriptional context of the naive and inflamed esophagus, enabling us to define disease-associated changes in immune and structural compartments.

### Th2 cells are the predominant cells expressing type 2 cytokines in experimental EoE

To characterize the lymphoid compartment, we re-clustered lymphocytes from naive and OXA-challenged esophagi. This identified CD4⁺ and CD8⁺ T-cell subsets, NK cells, γδT17 cells, ILC2s, B cells, and plasma cells (Fig. 2A-B). Th2 cells expressed *Gata3* and *Il1rl1* together with *Il4*, *Il13*, and *Calca*, whereas regulatory T cells (Tregs) expressed *Foxp3*, *Ikzf2*, and *Il2ra*. CD8⁺ T cells included circulating-like, cycling, and resident memory-like populations, with the latter expressing *Itgae*, *Itga1*, and *Cxcr6*. Cycling T cells expressed *Mki67* and *Top2a* (Fig. 2A-B).

**Figure 2.**
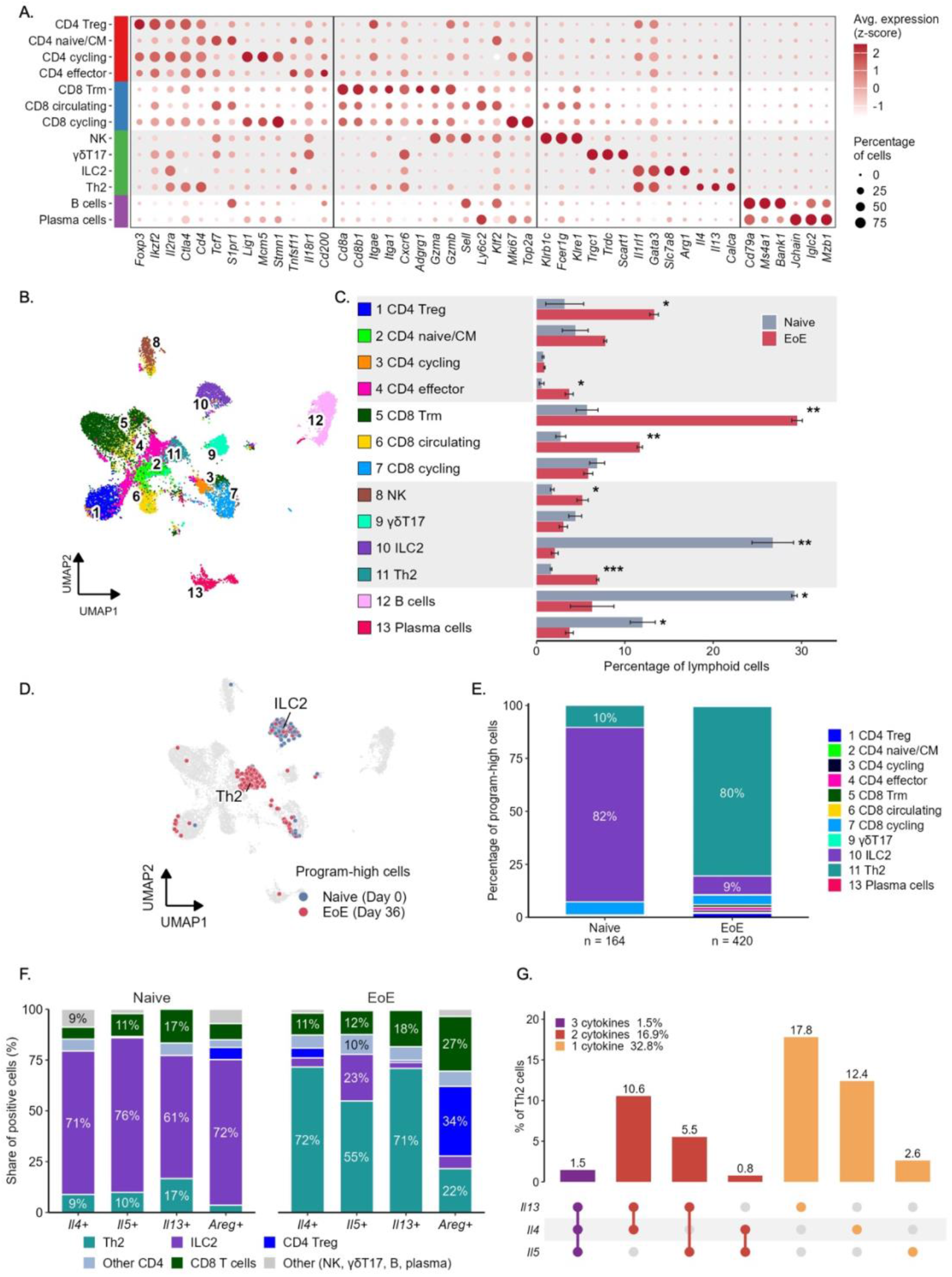
Th2 cells replace ILC2s as the main lymphoid source of type 2 cytokines in experimental EoE. Lymphoid cells were re-clustered and the expression of marker genes is shown (A). Dot color represents the average expression (z-score) and dot size represents the percentage of cells expressing each gene (A). UMAP representation of lymphoid populations is presented (B), and the percentage of each population among lymphoid cells in naive and EoE mice was determined (C). Type 2 program-high cells (top 10% of a score of *Il13*, *Il4*, *Il5*, *Areg*, *Gata3* and *Il1rl1*, with at least one of *Il4*, *Il5* or *Il13* detected) are shown on the lymphoid UMAP (D), and their composition in naive and EoE mice is presented (E). The contribution of each population to *Il4*⁺, *Il5*⁺, *Il13*⁺ and *Areg*⁺ lymphoid cells was determined (F). Co-expression of *Il13*, *Il4* and *Il5* in Th2 cells of EoE mice is shown as an UpSet plot (G); In (C), data are presented as mean ± SEM of two pooled samples per condition; *p < 0.05, p < 0.01,* p < 0.001 (two-tailed unpaired Student’s t-test).

Experimental EoE was associated with a marked change in the composition of the recovered lymphoid compartment. In naive mice, B cells and ILC2s accounted for ∼30% and 24-29% of lymphocytes, respectively, followed by plasma cells (Figure 2C). Consistent with previous studies in humans^1,2^, following OXA challenge, resident memory-like CD8⁺ T cells became the largest lymphoid population displaying a 5-fold increase (∼6% naive mice vs. ∼30% in EoE) and circulating CD8⁺ T cells increased ∼4-fold (from ∼3% to ∼12%, Figure 2C). Tregs and Th2 cells also expanded by ∼4.5-fold and ∼4.2-fold, respectively (increasing from ∼3% to ∼14% and from ∼1.6% to ∼7%, respectively).

Given the increased representation of Th2 cells, we next examined the distribution of type 2-associated gene expression among lymphocytes. Individual cells were scored using *Il4*, *Il5*, *Il13*, *Areg*, *Gata3*, and *Il1rl1*. This gene program was specifically selected to capture complementary features of type 2 immunity, including effector cytokines (*Il4*, *Il5*, and *Il13*), transcriptional identity (*Gata3*), the capacity to sense IL-33 (*Il1rl1*), and type 2 tissue growth- and repair-associated molecules (*Areg*)^18^. We selected cells with a score in the highest decile and detectable expression of at least one of the three cytokine transcripts, identifying 164 cells in naive mice and 420 in OXA-challenged mice. ILC2s accounted for 82% of these cells in naive mice, whereas Th2 cells accounted for 80% following OXA challenge (Fig. 2D-E). Analysis of individual cytokines showed a similar pattern. In naive mice, ILC2s accounted for 71%, 76%, and 61% of *Il4*-, *Il5*-, and *Il13*-expressing lymphocytes, respectively. Following OXA challenge, Th2 cells accounted for 72%, 55%, and 71%, respectively, while ILC2s retained a substantial contribution to *Il5* expression (23%; Fig. 2F). Within the Th2 population, approximately 33% expressed one of these cytokine genes, 17% expressed two, and only 1.5% expressed all three. The most frequent patterns were *Il13* alone and co-expression of *Il13* with *Il4* (Fig. 2G). Interestingly, *Areg* expression was more broadly distributed. Tregs, CD8⁺ T cells, and Th2 cells accounted for 34%, 27%, and 22% of *Areg*-expressing lymphocytes in OXA-challenged mice, respectively (Fig. 2F).

To examine convergence with human disease, we applied the corresponding scoring approach to a published human esophageal dataset (SCP1242)^12^. Th2 cells accounted for 57%, 78%, and 85% of high-scoring lymphocytes in control individuals, patients in remission, and patients with active EoE, respectively, whereas ILC2s accounted for 3%, 8%, and 3% (Fig. S1A-B). Thus, Th2 predominance was evident even in human controls. Comparison of curated gene categories further supported shared features of mouse and human Th2 cells, including type 2 cytokines, Th2 identity, effector mediators, and activation-associated genes (Fig. S1C).

Together, these findings identify a change in the predominant type 2 cytokine-expressing lymphocytes in experimental EoE and further reveal concordance between mouse and human Th2 cells in EoE.

### Th2 cells adopt a tissue-adapted effector state that extends beyond cytokine production

Since Th2 cells were the predominant source of type 2 cytokines in experimental EoE, we next sought to define their mediator repertoire. To this end, we compared the transcriptional profile of Th2 cells with that of all other lymphocytes recovered from OXA-challenged mice and, separately, with that of effector CD4⁺ T cells, which allowed us to distinguish Th2-specific features from those shared with activated CD4⁺ T cells (Fig. 3A). Beyond their annotating transcripts (*Il4*, *Il5*, *Il13*, *Gata3* and *Il1rl1*), Th2 cells were enriched for the secreted mediators *Calca*, *Il10*, *Areg* and *Fgl2*, for the surface receptors *Cysltr1*, *Nmur1*, *Cd200r1* and *Klrg1*, for the transcriptional regulators *Maf*, *Irf4* and *Pparg*, and for the metalloproteinase *Adam8*. Notably, *Calca*, which encodes calcitonin gene-related peptide (CGRP)^19^, and *Foxb1* were among the most highly enriched transcripts in both comparisons.

**Figure 3.**
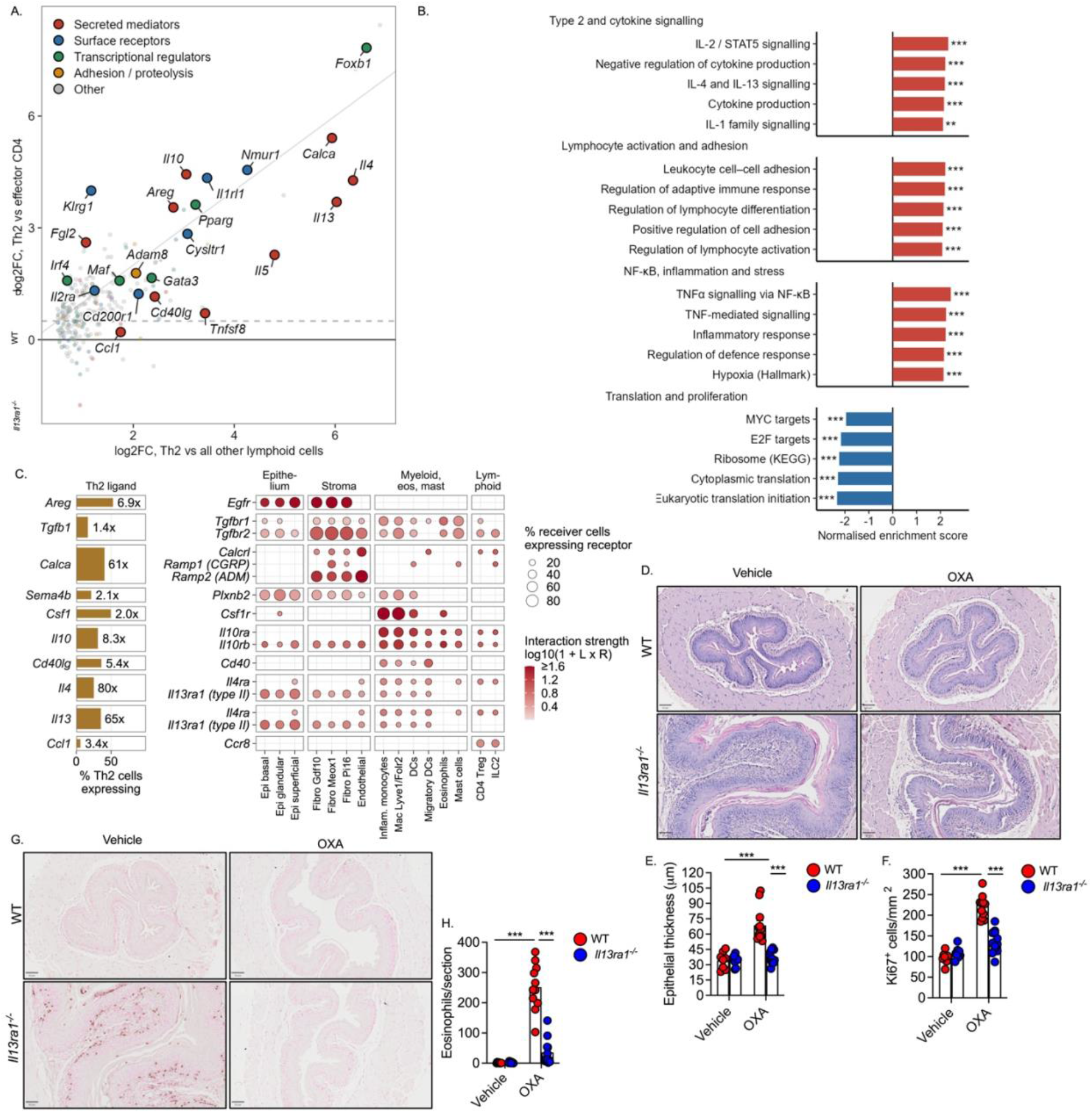
Th2 cell-derived mediators act on IL-13Rα1-expressing structural cells to promote experimental EoE. Genes enriched in Th2 cells of mice with EoE were compared with all other lymphoid cells (x-axis) and with effector CD4⁺ T cells (y-axis) (A). Gene set enrichment analysis (GSEA) of Th2 cells compared with all other lymphoid cells at day 36 is presented (B). Ligands expressed by Th2 cells (percentage of Th2 cells and fold enrichment over other lymphoid cells) and the expression of their receptors in epithelial, stromal, myeloid and lymphoid cells are shown (C); dot size represents the percentage of receiving cells expressing the receptor and dot color the interaction strength (C). Experimental EoE was induced in WT and *Il13ra1^⁻/⁻^* mice using OXA; control mice received vehicle. On day 36, the mice were euthanized, and esophageal tissues were fixed, paraffin embedded, and slides were generated. H&E-stained stained slides were analyzed (D) and epithelial thickness was determined (E). Epithelial cell proliferation was determined using anti-Ki67 staining (F). The slides were stained with anti-major basic protein [MBP (G)] and esophageal eosinophils were quantified (H). scale bars, 50 µm. In (B), *adjusted p < 0.01*, *adjusted p < 0.001. In (E, F, H), each dot represents one mouse; data are presented as mean ± SEM with 8–10 mice per group;* p < 0.001 (two-way ANOVA).

To determine which cellular programs underlie this mediator profile, we performed ranked pathway analysis (Fig. 3B). Th2 cells were enriched for cytokine production and signaling (e.g., IL-2/STAT5, IL-4 and IL-13, and IL-1 family signaling), for lymphocyte activation and adhesion, and for TNFα/NF-κB signaling. Negative regulation of cytokine production was also enriched, consistent with the concurrent enrichment of regulatory-associated transcripts (e.g., *Cd200r1*, *Klrg1* and *Il10*, Fig. 3A). In contrast, translation- and proliferation-associated pathways including MYC and E2F targets, ribosomal genes and translation initiation were depleted. Thus, esophageal Th2 cells display a chronically activated, non-proliferating effector state in which cytokine production is accompanied by intrinsic regulatory programs.

To identify potential links between the mediator repertoire of Th2 cells and target cells in the esophagus, ligands expressed by Th2 cells were paired with receptor expression in epithelial, fibroblast and endothelial populations, as well as in selected immune populations (Fig. 3C). *Il4* and *Il13* were the most highly Th2-enriched ligands (80- and 65-fold, respectively), followed by *Calca* (61-fold), *Il10* (8.3-fold), *Areg* (6.9-fold) and *Cd40lg* (5.4-fold). *Il4* and *Il13* were predicted to engage *Il13ra1*, which was expressed by epithelial cells and fibroblasts, whereas *Il4ra* was expressed mainly by myeloid, mast and lymphoid populations. *Areg*, *Sema4b* and *Tgfb1* were paired with *Egfr*, *Plxnb2* and *Tgfbr1*/*Tgfbr2* on these same structural populations. *Csf1*, *Il10* and *Cd40lg* were predicted to interact with myeloid cells through *Csf1r*, *Il10ra*-*Il10rb* and *Cd40*.

Consistent with the predicted interaction between IL-4 and IL-13 with IL-13Rα1, OXA-challenged *Il13ra1*^-/-^ mice were markedly protected from EoE^16^, displaying decreased basal cell proliferation and epithelial thickening (Fig. 3D-F) as well as decreased accumulation of MBP⁺ eosinophils (Fig. 3G-H). Together, these findings identify esophageal Th2 cells as a source of mediators that extend beyond type 2 cytokines and establish IL-13Rα1 on structural cells as a required target of this response.

### Lymphoid populations acquire a shared interferon program in experimental EoE

To determine whether the effector state identified in Th2 cells is confined to this population or is shared more broadly within the lymphoid compartment, we performed ranked pathway analysis on ILC2s, Tregs and resident memory-like CD8⁺ T cells from OXA-challenged compared with naive mice (Fig. S2). The interferon-γ response was enriched in all three populations, together with the interferon-α response in ILC2s and Tregs, TNF/NF-κB signaling in Tregs, and cytotoxic effector activity in resident memory-like CD8⁺ T cells (Fig. S2A-F). Notably, the interferon response in ILC2s was accompanied by a loss of their type 2 program, since *Il5*, *Il13*, *Il1rl1* and *Gata3* were lower following OXA challenge and only *Areg* was consistently higher (Fig. S2B). These data identify a shared interferon-associated program, indicating that the lymphoid response in experimental EoE extends beyond type 2 immunity.

### Experimental EoE alters myeloid composition and reveals heterogeneous macrophage states

To characterize the myeloid response in experimental EoE, we re-clustered 18,507 recovered myeloid cells, which resolved 16 populations comprising classical monocytes, neutrophils, eosinophils, five macrophage states, four dendritic cell states, Langerhans cells, cycling myeloid cells, and connective tissue-type and mucosal-type mast cells (Fig. 4A, 4B). Macrophage heterogeneity included populations annotated by *Lyve1* and *Folr2*, *C1q* and *Cx3cr1*, or *Retnla* and *Lpl* expression, as well as inflammatory and interferon-associated *Itgax*⁺*Cxcl9*⁺ states.

**Figure 4.**
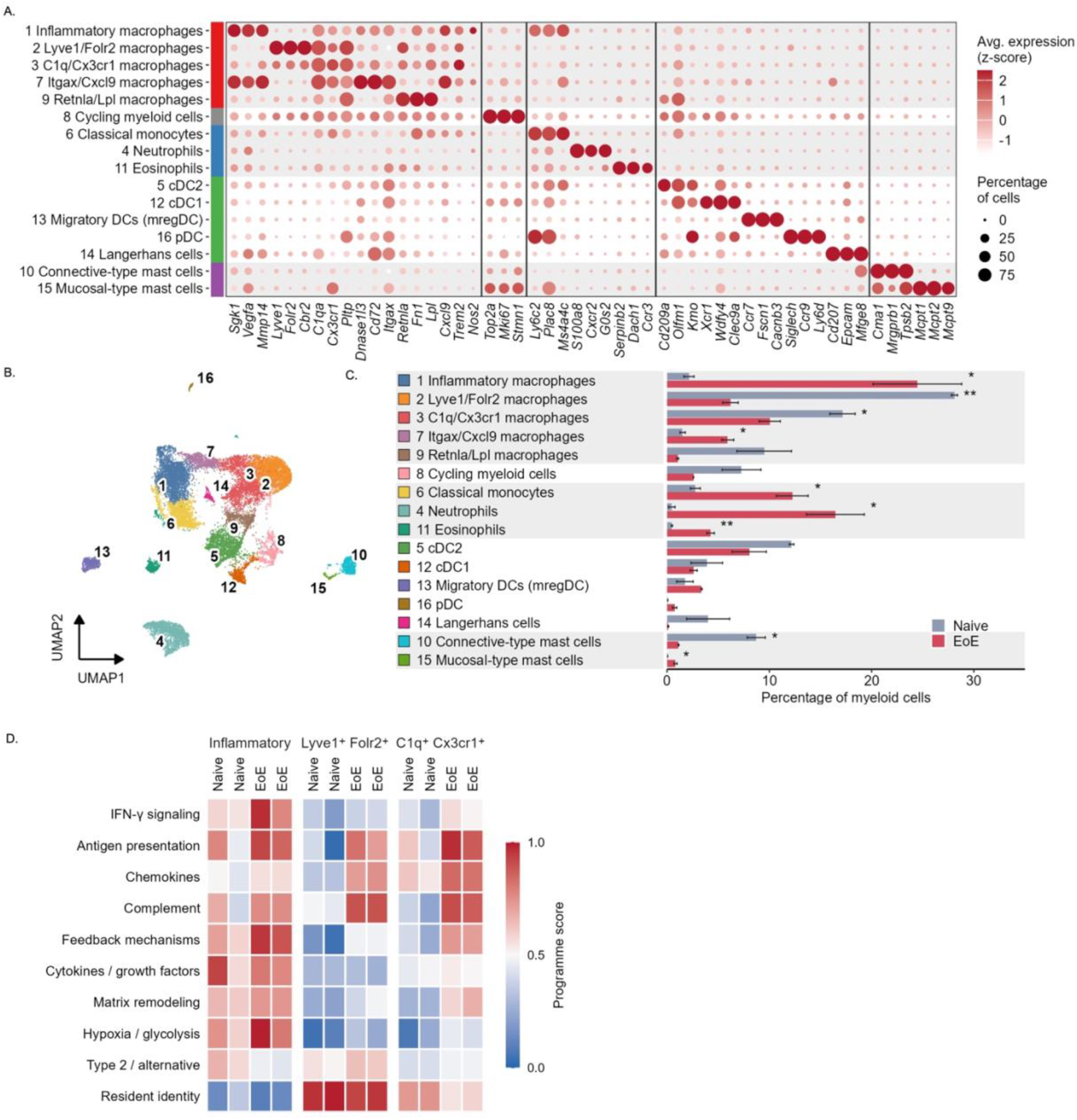
Inflammatory macrophages replace resident macrophage populations in experimental EoE. Myeloid cells were clustered and the expression of marker genes is shown (A). Dot color represents the average expression (z-score) and dot size the percentage of cells expressing each gene (A). UMAP representation of myeloid clusters is presented (B), and the percentage of each cluster among myeloid cells in naive and EoE mice was determined (C). Functional programs of inflammatory (D-left panel), Lyve1⁺Folr2⁺ (D-middle panel) and C1q⁺Cx3cr1⁺ (D-right panel) macrophages are shown for each sample (D). For each gene, pseudobulk expression was scaled from 0 to 1 across the 12 population-by-sample values, and the program score presented as the mean over the program genes listed in Fig. S3. In (C), data are presented as mean ± SEM of two pooled samples per condition; *p < 0.05, p < 0.01,* p < 0.001 (two-tailed unpaired Student’s t-test).

The composition of the myeloid compartment differed markedly between naive and OXA-challenged mice (Fig. 4C). In naive mice, the compartment was predominantly composed of resident macrophage states including *Lyve1*⁺*Folr2*⁺ (∼28%) and *C1q*⁺*Cx3cr1*⁺ (∼17%) macrophages. Following OXA challenge, this resident network was largely replaced by recruited and inflammatory populations. In fact, inflammatory macrophages expanded from 2% to 24% of myeloid cells and became the dominant population. This was accompanied by the infiltration of neutrophils, classical monocytes and eosinophils, which were nearly absent at baseline. An increase in interferon-associated *Itgax*⁺*Cxcl9*⁺ macrophages and migratory DCs was observed as well (Fig. 4C). Mast cells underwent a notable switch, with loss of connective tissue-type mast cells and emergence of mucosal-type mast cells following OXA challenge (Fig. 4C). Thus, experimental EoE was associated with broad alterations of the myeloid compartment.

The expansion of inflammatory macrophages raised the question of whether the resident macrophage populations that persisted in the inflamed esophagus also altered their transcriptional state. To address this, we scored ten pre-specified gene programs in the three most abundant macrophage populations and compared them between naive and OXA-challenged mice (Fig. 4D; individual genes in Fig. S3). Following OXA challenge, interferon-γ signaling, antigen presentation, chemokine and complement programs were elevated in all three populations (Fig. 4D). Inflammatory macrophages scored highest for interferon-γ signaling, feedback mechanisms, hypoxia and glycolysis, consistent with increased expression of *Il18bp*, *Il1rn* and *Havcr2* (Fig. S3). The resident macrophage populations were reprogrammed in this inflammatory environment. Chemokine and complement programs were upregulated in *Lyve1*⁺*Folr2*⁺ and *C1q*⁺*Cx3cr1*⁺ macrophages (Fig. 4D), with induction of the T cell-attracting chemokines *Ccl8* and *Cxcl9* (Fig. S3). Finally, despite acquiring these shared programs, the two resident populations differed in the fate of their baseline state, since *C1q*⁺*Cx3cr1*⁺ macrophages lost part of their resident identity whereas *Lyve1*⁺*Folr2*⁺ macrophages retained it (Fig. 4D).

These data identify an interferon-associated, antigen-presenting and chemokine-producing macrophage state as a dominant myeloid program in experimental EoE, which mirrors the interferon-enriched program observed in the lymphoid compartment.

### Esophageal eosinophils integrate epithelial, stromal and lymphoid signals in EoE

Esophageal eosinophilia is the histological hallmark of EoE and serves as a diagnostic and monitoring criterion^20, 21^. Yet the activities of eosinophils within the inflamed esophagus remain to be defined. To define their transcriptional profile, we compared eosinophil gene expression with that of all other myeloid cells recovered from OXA-challenged mice. This analysis identified 3,423 differentially expressed genes (adjusted p < 0.05), of which 634 were higher in eosinophils (Fig. S4). The enriched genes comprised programs of mediator secretion (*Serpinb2*, *F5*, *Retnlg*), lipid-mediator synthesis (*Alox15*, *Pla2g7*, *Pla2g4a* and *Lta4h*) and survival (*Morrbid*), rather than granule protein production (Fig. 5A). In contrast, the type 2 transcripts *Il4* and *Il1rl1* were detected only in a minority of eosinophils, and among all myeloid cells, were expressed predominantly by mast cells (Fig. 5A).

**Figure 5.**
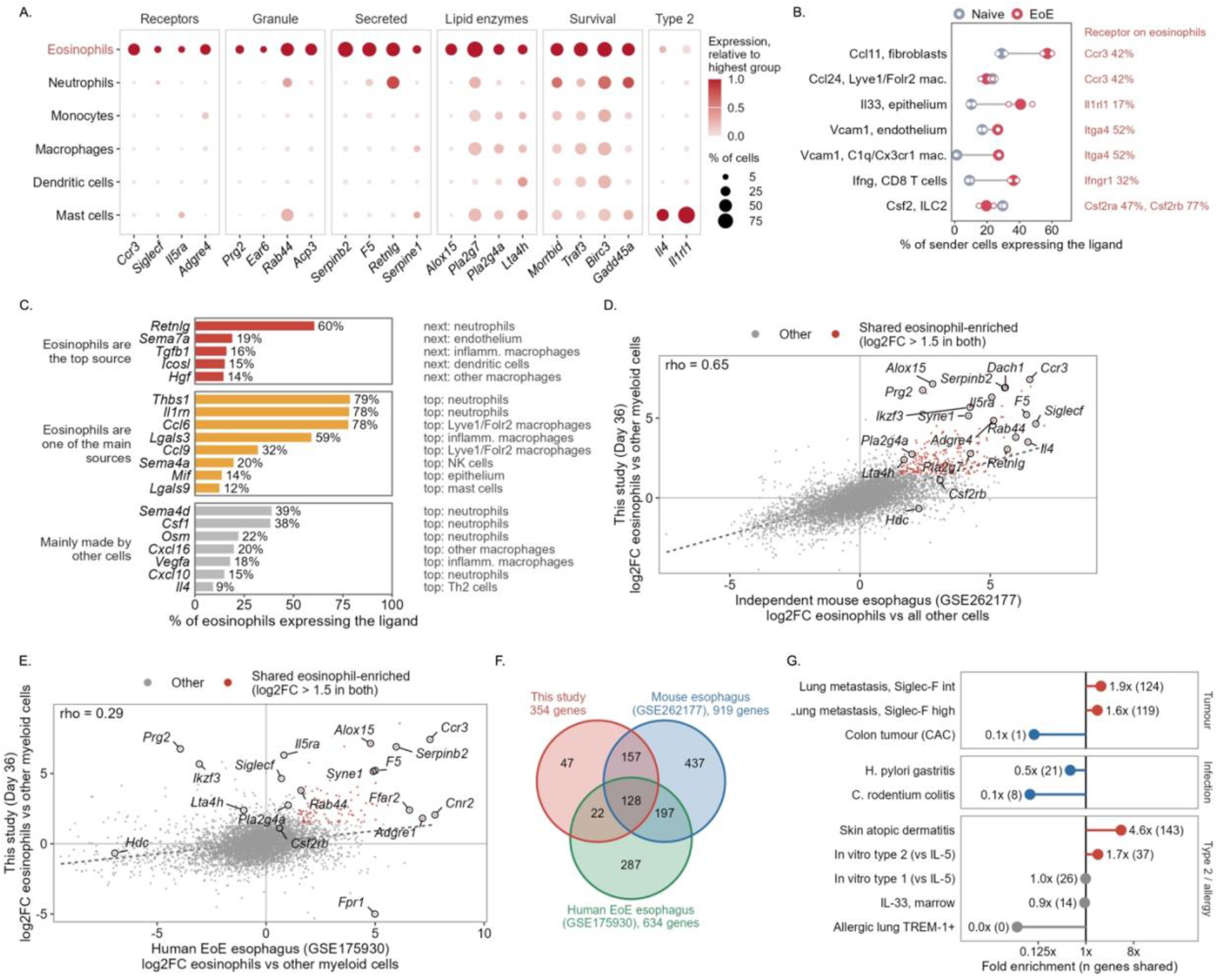
Esophageal eosinophils display a conserved tissue program in experimental EoE. Expression of receptors, granule proteins, secreted mediators, lipid enzymes, survival regulators and type 2 genes in myeloid groups of experimental EoE mice is shown (A). Dot color represents the expression relative to the highest group and dot size the percentage of cells expressing each gene (A). Ligands that can interact with eosinophils were paired with receptors detected in at least 10% of eosinophils. The percentage of sender cells expressing each ligand in naive and experimental EoE mice as well as the percentage of eosinophils expressing the receptor are shown (B). Ligands expressed by eosinophils, grouped by whether eosinophils are the top sources, one of the main or a minor source, are presented (C). Eosinophil-enriched genes (log2 fold change, eosinophils versus other myeloid cells) were compared with an independent mouse esophagus dataset (GSE262177; D) and with esophageal eosinophils from patients with active EoE (GSE175930; E). Red dots represent genes enriched more than 2.8-fold in both datasets and rho the Spearman correlation. The overlap between eosinophil-enriched genes of this study, the mouse esophagus dataset and human EoE eosinophils are shown (F). The overlap with genes induced in eosinophils in lung metastasis (GSE181491), colitis-associated colon tumor (GSE112922), *Helicobacter pylori* gastritis, *Citrobacter rodentium* colitis (GSE182001), atopic dermatitis (GSE224703), in vitro type 2 and type 1 conditions (GSE243369), IL-33-exposed bone marrow (GSE249011) and allergic lung TREM-1⁺ eosinophils (GSE246389) is presented as fold enrichment over chance, with the number of shared genes in brackets (G). In (B), open circles represent individual mice and filled circles the mean.

We next sought to define potential interactions between eosinophils and additional cells in the esophagus. To this end, ligands expressed by neighboring populations were paired with receptors detected in at least 10% of eosinophils (Fig. 5B). OXA challenge induced four ligands, each derived from a distinct cellular source. *Ccl11* from fibroblasts, *Il33* from the epithelium, *Vcam1* from endothelial cells and *C1q*⁺*Cx3cr1*⁺ macrophages, and *Ifng* from CD8⁺ T cells. The corresponding receptors *Ccr3*, *Il1rl1*, *Itga4* and *Ifngr1* were all expressed by eosinophils. Eosinophils themselves were the dominant source of *Retnlg*, *Sema7a*, *Tgfb1*, *Icosl* and *Hgf*, and were among the main sources of *Thbs1*, *Il1rn*, *Ccl6* and *Lgals3* (Fig. 5C). Expression of *Tgfb1*, *Thbs1* and *Hgf* suggests that eosinophils contribute to the tissue remodeling that accompanies EoE.

To determine whether this profile is shared by esophageal eosinophils more broadly, we compared gene-level contrasts with independent published datasets. Eosinophil enrichment in this study correlated with that in an independent murine esophageal EoE dataset (GSE262177^22^; Spearman ρ = 0.65; Fig. 5D) and, more weakly, with eosinophils from biopsies of patients with active EoE (GSE175930^23^; ρ = 0.29; Fig. 5E). Of the 354 eosinophil-enriched genes identified here, 150 were also enriched in human EoE eosinophils, representing 23% of the human eosinophil-enriched gene set (Fig. 5F). Comparison with eosinophil signatures from other disease contexts revealed the strongest overlap with skin eosinophils in atopic dermatitis^24^, whereas overlap with eosinophils from other diseased mucosal organs such as the lungs (GSE181491, GSE246389)^25, 26^, colitis-associated tumors (GSE112922)^27^, *C. rodentium* colitis and *H. pylori* gastritis (GSE182001)^28^, was below that expected by chance (Fig. 5G). The esophageal profile was not reproduced by cytokine polarization in vitro (GSE243369, GSE249011)^29^. Collectively, these data suggest that the esophageal eosinophil transcriptional profile in EoE is shaped by local tissue cues rather than by type 2 cytokines alone.

### Mucosal-type mast cells emerge in experimental EoE and express a distinct mediator repertoire

The finding that mast cells, rather than eosinophils, were the principal myeloid source of type 2 transcripts in the inflamed esophagus (Fig. 5A) prompted us to examine this compartment more closely. Re-clustering of esophageal mast cells distinguished connective tissue-type cells, defined by *Cma1*, *Mcpt4* and the *Mrgpr* receptors, from mucosal-type cells, defined by *Mcpt1* and *Mcpt2* (Fig. 6A). Mast cells in naive mice were almost exclusively connective tissue-type. Following OXA challenge, mucosal-type cells comprised approximately half of the compartment and half of these were cycling (*Mki67*, *Top2a*; Fig. 6A). This shift was confirmed at the tissue level, since MCPT1⁺ cells were rare in vehicle-treated mice and readily detected in OXA-challenged esophagi (Fig. 6B).

**Figure 6.**
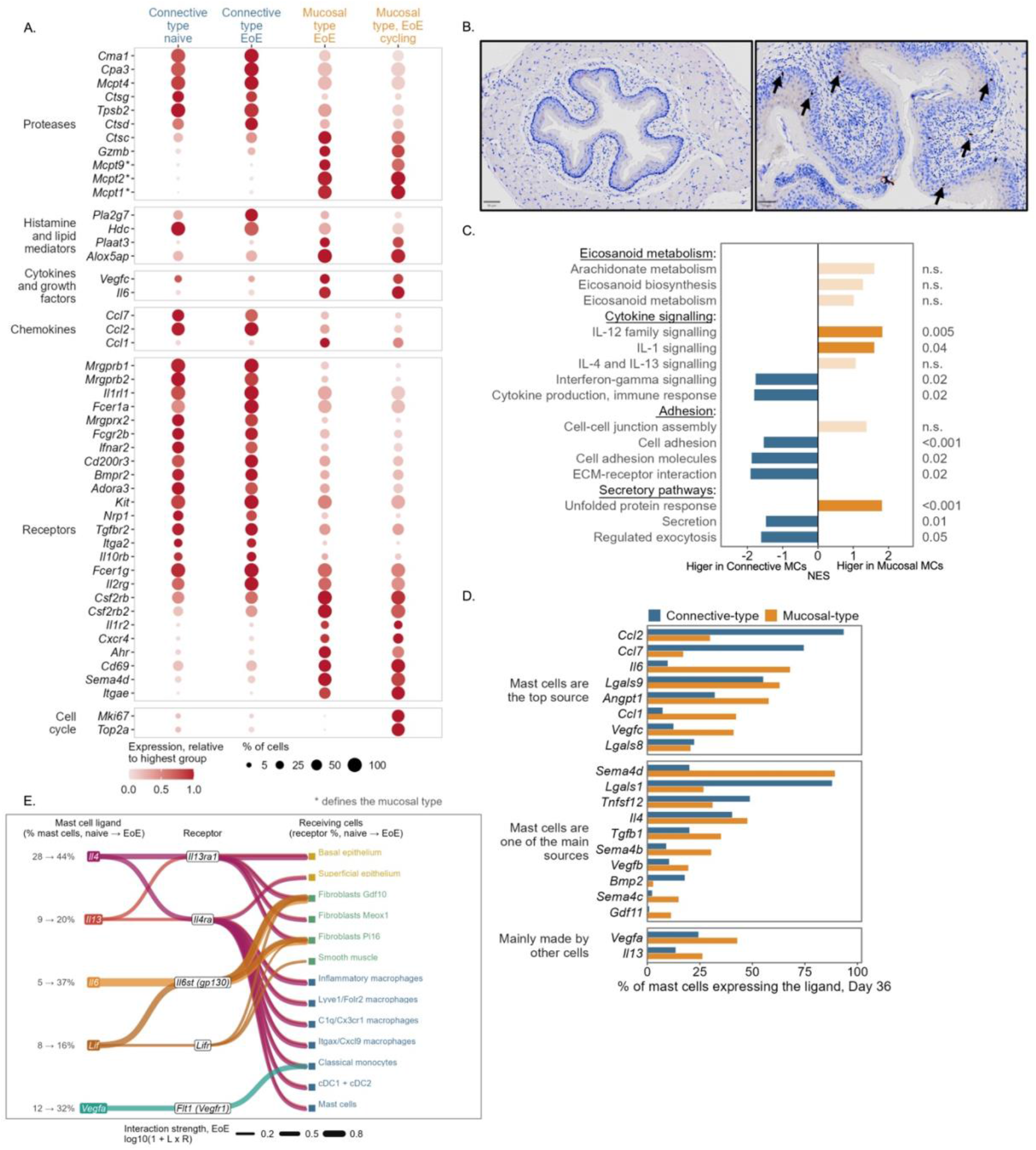
Experimental EoE is characterized by the emergence of mucosal-type mast cells. Expression of proteases, histamine and lipid mediator genes, cytokines, chemokines, receptors and cell-cycle genes in connective-type mast cells of naive and EoE mice and in non-cycling and cycling mucosal-type mast cells of EoE mice is shown (A). Dot color represents the expression relative to the highest group and dot size the percentage of cells expressing each gene (A). Esophageal tissues of naive and OXA-challenged mice were stained with anti-MCPT1. A representative photomicrographs are shown and arrows indicate MCPT1⁺ cells (B, scale bars, 50 µm). GSEA of mucosal-type compared with connective-type mast cells is presented, with adjusted p values (C). Ligands expressed by connective-type and mucosal-type mast cells in EoE mice are shown (D). Interactions in which both the mast cell ligand and the receptor on the receiving population increased in EoE (Wilcoxon test, adjusted p < 0.05, ≥ 1.2-fold; receptor detected in ≥ 10% of receiving cells) are presented (E); numbers represent the percentage of mast cells expressing each ligand (naive → EoE) and line width the interaction strength.

The two mast cell types differed in 963 genes, of which 511 were higher in mucosal-type and 452 in connective tissue-type cells (Fig. S5A). Mucosal-type cells were distinguished by *Il6*, *Ccl1* and *Vegfc*, by the 5-lipoxygenase pathway components *Alox5ap* and *Ptgs2*, and by enrichment of IL-1 and IL-12 family signaling and the unfolded protein response (Fig. 6A, 6C). Connective tissue-type cells expressed the monocyte-attracting chemokines *Ccl2* and *Ccl7* and were enriched for cell adhesion, extracellular matrix-receptor interaction and regulated exocytosis (Fig. 6A, 6C).

To relate mast cell-expressed mediators to the tissue changes of EoE, we compared mast cell ligands with their expression in all other esophageal populations (Fig. 6D). Mucosal-type mast cells were the dominant source of *Il6*, *Ccl1*, *Vegfc* and *Angpt1*, and both mast cell types were a major source of *Il4*, second only to Th2 cells. We next examined ligand-receptor pairs in which both the mast cell ligand and its receptor increased following challenge (Fig. 6E, Fig. S5C). Mast cell expression of *Il4*, *Il13*, *Il6* and *Lif* was induced, and the corresponding receptors were concurrently upregulated on the structural compartments of the esophagus. Consistent with the dominant role of IL-13Rα1, *Il4* and *Il13* were paired with *Il13ra1*, which increased on basal epithelial cells and on *Gdf10*⁺, *Meox1*⁺ and *Pi16*⁺ fibroblasts. Mast cells also received signals from these same compartments, since epithelial *Il33*, fibroblast- and endothelium-derived *Cxcl12*, and CD8⁺ T cell-derived *Ifng* were paired with *Il1rl1*, *Cxcr4* and *Ifngr1* on mast cells, respectively (Fig. S5B).

To determine whether these programs are conserved in mast cells from human disease, we compared gene-level contrasts using a published dataset (GSE201153^30^; Fig. S5D). The overall correlation was modest (Spearman ρ = 0.28 over 3,622 orthologous genes), and the transcripts that were mast cell-specific in both species comprised the lineage-determining transcription factors, the granule proteases and the histamine-handling machinery. Thus, the lineage identity of esophageal mast cells is conserved between experimental and human EoE, whereas their effector repertoires diverge.

### Experimental EoE is associated with epithelial stress and inflammatory responses in esophageal fibroblasts

EoE is characterized by barrier disruption and fibrosis^2, 31^. Thus, we aimed to determine how experimental EoE alters the gene programs of non-hematopoietic cells in the esophagus. Following OXA challenge, the keratin gradient of the epithelium was disrupted. Basal cells displayed reduced *Krt15*, whereas the basal keratin *Krt14* was detected in a large proportion of superficial cells, and both layers induced the stress-associated keratins *Krt6a* and *Krt16* (Fig. 7A). These changes were confirmed at the protein level by KRT6 and KRT14 staining (Fig. 7B). Superficial cells further displayed higher *Sprr1a* and reduced *Dsg1a*, consistent with the barrier impairment that characterizes human EoE. As previously shown, *Mki67* was induced in both epithelial layers (Fig. 7A). Beyond this injury response, epithelial cells displayed a mixed response characterized by an interferon signature (*Stat1*, *Cxcl9*, *H2-K1*, *B2m*), a type 2 signature (*Alox15*, *Ccl11* and *Chil4*), an alarmin signature (*Il33*, *Il36g* and *Tslp*) and induction of chemokines (*Ccl2* and *Ccl8*) (Fig. 7A). Additional interactions included *Cxcl16* that could potentially recruit CD8⁺ T cells^32^. Epithelial cells also displayed increased *Il13ra1* and *Ifngr1*, indicating that they are positioned to respond to both arms of this response.

**Figure 7.**
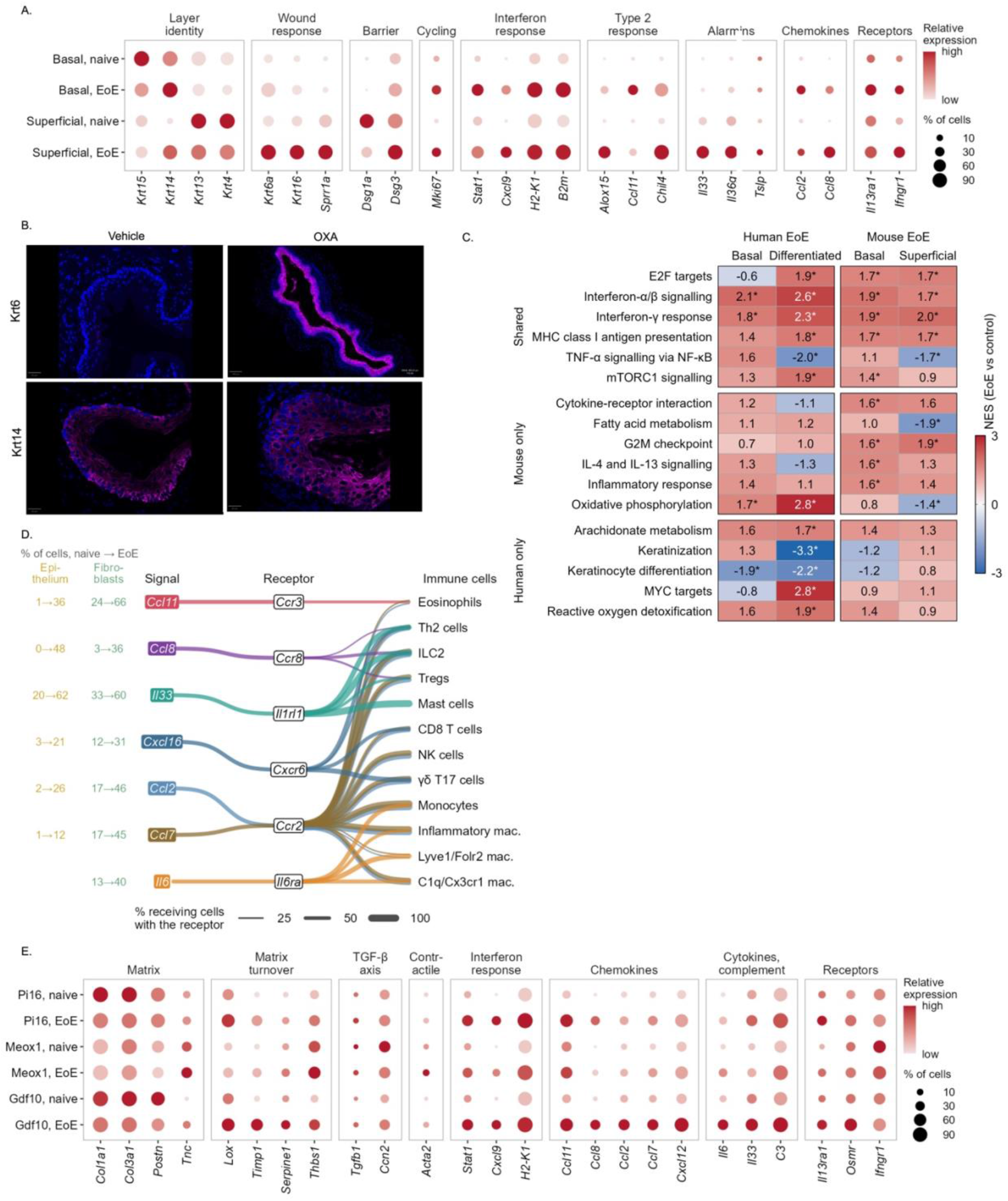
Structural cells mount interferon and inflammatory responses in experimental EoE. Expression of layer identity, wound response, barrier, proliferation, interferon response, type 2 response, alarmin, chemokine and cytokine receptor genes in basal and superficial epithelial cells of naive and EoE mice is shown (A). Dot color represents the expression relative to the highest group and dot size the percentage of cells expressing each gene (A). Esophageal tissues from vehicle- and OXA-treated mice were stained with anti-KRT6 or anti-KRT14 (magenta) and DAPI (blue), representative photomicrographs are shown (B, scale bars, 50 µm). GSEA of epithelial cells from patients with active EoE (GSE201153) and from experimental EoE is presented as normalized enrichment scores of EoE versus control (C). Pathways are grouped as shared, mouse or human only ( adjusted p < 0.05). Ligands that increased in epithelial cells or fibroblasts were paired with receptors on immune populations (D). Numbers represent the percentage of epithelial cells and fibroblasts expressing each ligand (naive → EoE) and line width the percentage of receiving cells expressing the receptor. Expression of matrix, matrix turnover, TGF-β axis, contractile, interferon response, chemokine, cytokine, complement and receptor genes in Pi16, Meox1 and Gdf10 fibroblasts of naive and EoE mice is shown (E). Dot color represents the expression relative to the highest group and dot size the percentage of cells expressing each gene.

To determine whether these transcriptional responses reflect those observed in patients, we compared gene set enrichment between the mouse epithelial layers and the epithelium of patients with active EoE (GSE201153^30^, Fig. 7C). Proliferation, interferon-α/β signaling, the interferon-γ response and MHC class I antigen presentation were enriched in both species. Interestingly, despite elevation in human basal cells, IL-4 and IL-13 signaling reached significance only in mouse basal cells (Fig. 7C). Furthermore, the loss of keratinization that characterized patient epithelium was not reproduced in experimental EoE. Therefore, experimental EoE recapitulates the proliferative and interferon-associated epithelial response of human EoE, but only part of its IL-13-dependent signature and not its loss of terminal differentiation.

We next compared esophageal fibroblasts between naive and OXA-challenged mice (Fig. 7E). *Col1a1* and *Col3a1* were expressed by virtually every fibroblast in both groups and were not further induced, whereas genes regulating matrix turnover were increased, including *Timp1*. *Lox*, *Serpine1* and *Thbs1* were predominantly increased in *Pi16*- and *Gdf10*-expressing fibroblasts (Fig. 7E). *Meox1*-expressing fibroblasts additionally induced *Tnc* and *Acta2*, consistent with a myofibroblast-like state. Importantly, all three populations acquired inflammatory programs, with induction of an interferon response (*Stat1*, *Cxcl9*, *H2-K1*) and increased *C3*, *Il6*, *Il33* and *Il13ra1*. In addition, *Gdf10*-expressing fibroblasts induced a broad chemokine response, with induction of *Ccl2*, *Ccl7* and *Cxcl12*. *Ccl11*, a key eosinophil-attracting chemokine, was detected in most *Pi16*- and *Gdf10*-expressing fibroblasts.

To determine which immune populations could respond to these structural cell-derived signals, we matched ligands that increased in epithelial cells or fibroblasts with receptor expression in immune populations (Fig. 7D). *Ccl11* was predicted to signal to eosinophils through *Ccr3*. *Ccl8* was predicted to signal to Th2 cells, ILC2s and Tregs through *Ccr8*. Although *Il33* had a limited role in experimental EoE, it was predicted to signal to multiple cells including Th2 cells, ILC2s, Tregs and mast cells through *Il1rl1*. These data position the esophageal epithelium and fibroblasts as both targets of IL-13 and IFN-γ and sources of the chemokines and alarmins that recruit and sustain the immune compartment of experimental EoE.

## Discussion

The advent of biological therapies in allergic diseases has increasingly revealed the dependence of disease on individual cytokines. In EoE, clinical trials and studies using experimental models have demonstrated important roles for IL-4, IL-13, IL-4Rα, TSLP and IL-33^5, 6, 16, 33, 34, 35, 36, 37^. Herein, we defined the immune and structural cell landscape of experimental EoE and analyzed which cells may respond and/or produce this cytokine profile. In addition, we compared it with human disease. Our analyses identify four principal findings. First, Th2 cells replace ILC2s as the predominant source of type 2 cytokines in the inflamed esophagus. Second, the mast cell compartment switches from a connective tissue-type population to a largely mucosal-type and partly cycling state. Third, esophageal eosinophils adopt a signature that resembles that of skin eosinophils in atopic dermatitis rather than that of eosinophils from other gastrointestinal tissues. Fourth, an interferon-associated program accompanies the type 2 response in lymphoid, myeloid and structural compartments, and its epithelial component is conserved in human EoE.

Our study identifies Th2 cells as the predominant source of type 2 cytokines in the inflamed esophagus, which is consistent with human studies identifying highly cytokine-expressing tissue Th2 cells in active EoE^8, 23, 38, 39, 40, 41^. Notably, this predominance was not evident at baseline, since ILC2s accounted for most type 2 cytokine-expressing lymphocytes in naive mice, whereas Th2 cells already predominated in healthy human esophagus. This likely reflects the composition of the naive murine esophagus, in which KLRG1⁺ ILC2s are abundant under homeostatic conditions^42^. A sequential relationship between ILC2s and Th2 cells has been established in other allergic tissues^43^. ILC2-derived IL-13 licenses dendritic cells to produce CCL17 and is required for the recruitment of memory Th2 cells to the lung and skin following allergen challenge^44, 45^. Our data extend this relationship to the esophagus. They further indicate that this shift is not explained by Th2 cell expansion alone, since ILC2s concurrently downregulated *Il5*, *Il13*, *Il1rl1* and *Gata3* while acquiring an interferon-dominated program. Interestingly, *Areg* was the only type 2-associated transcript that ILC2s retained. This is of specific interest since esophageal ILC2s were recently shown to drive basal cell hyperproliferation and epithelial thickening through Areg-EGFR signaling independently of IL-5 and eosinophils^42^. Furthermore, inhibition of Areg and EGFR attenuated experimental EoE^42^. Thus, the reduced ILC2 contribution to cytokine expression does not indicate functional dispensability but rather suggests that their principal output in established disease is tissue remodeling rather than cytokine production. Since Tregs and CD8⁺ T cells were the other main *Areg*-expressing populations in our dataset, future studies determining the relative contribution of each *Areg* source to epithelial remodeling should be conducted.

The mast cell compartment underwent a parallel transition. At baseline, esophageal mast cells were almost exclusively connective tissue-type. Following challenge, mucosal-type cells comprised approximately half of the compartment, and half of these were cycling. Increased MCPT1⁺ cells in EoE were independently confirmed by immunohistochemistry. These findings correspond to human studies identifying heterogeneous esophageal mast cell populations in active EoE, including a transient proinflammatory subset that is a prominent source of IL-13^10^. This is clinically relevant since mast cell numbers and degranulation correlate with dysphagia in patients, and since elevated expression of mast cell CPA3 and chymase is associated with a steroid-resistant subtype of EoE^46, 47^. In our dataset, mucosal-type mast cells were a major source of *Il4* and engaged *Il13ra1* on epithelial cells and fibroblasts. Importantly, TSLP induces IL-13 secretion from mucosal but not connective tissue-type mast cells^17, 48^. Thus, the population that emerges in experimental EoE is the one that is responsive to TSLP. Nonetheless, these analyses should be interpreted with caution. Mucosal-type cells were nearly absent from the naive mouse esophagus whereas both protease-defined subsets are present in healthy human tissue. Furthermore, cross-species comparison indicated that while mast cell lineage identity is conserved between experimental and human EoE, their effector repertoires diverge. Therefore, the contribution of mast cells to EoE should be defined.

Recent studies, including our own, indicate that eosinophils are not required for the tissue pathology of EoE^7, 17^. Despite this, single-cell analysis of these cells is informative, since it provides insight into the environmental cues that shape their tissue adaptation^49, 50^. Esophageal eosinophils displayed a transcriptional signature that overlapped most strongly with that of skin eosinophils in atopic dermatitis, whereas overlap with gastrointestinal eosinophils fell below that expected by chance. The esophagus and the skin are lined by stratified squamous epithelium, whereas the stomach, intestine and colon are lined by columnar epithelium^51, 52^. Consequently, this pattern suggests that epithelial type rather than anatomical proximity dictates the transcriptional state of eosinophils. An alternative explanation is that this similarity reflects a shared sensitization protocol rather than a shared epithelium. We consider this unlikely, since we recently demonstrated that identical oxazolone challenge elicits markedly distinct transcriptional programs in the esophagus and the stomach^53^. In addition, the esophageal eosinophil profile was not reproduced by cytokine polarization in vitro. We therefore propose that the epithelium is a principal determinant of eosinophil state in allergic inflammation. This proposal is testable, since it predicts that eosinophils recruited to the stomach in experimental eosinophilic gastritis will resemble gastric rather than esophageal eosinophils. Future studies comparing eosinophils recovered from squamous and columnar mucosae in the same animal will address these questions.

Our analyses demonstrate that resident memory-like CD8⁺ T cells constituted the largest lymphoid population at the disease endpoint. These cells displayed the most pronounced effector program of any lymphoid population, comprising cytotoxic and interferon-associated genes together with depletion of proliferation signatures. Of note, interferon-associated programs were not confined to this population. They also characterized ILC2s, Tregs, resident macrophages, mast cells, epithelial cells and fibroblasts. Furthermore, the IFN-γ signature of the epithelial component was shared with human EoE. These observations converge with emerging human data suggesting Involvement of IFN-γ in EoE^54, 55^. Interferon-stimulated genes are most prominently induced in the epithelium of patients with active EoE, where they correlate with eosinophil counts and endoscopic severity^56^. In addition, IFNG is expressed by esophageal CD69⁺CD8⁺ T cells, and IFN-γ impairs epithelial barrier formation in esophageal organoids^56^. This interferon program operates within a disease that is highly dependent on type 2 signaling. Indeed, dupilumab is clinically effective in EoE^5, 6, 33^, and the phase 3 CROSSING trial recently demonstrated that TSLP blockade with tezepelumab induces histologic remission, improves dysphagia and reduces endoscopic severity in patients with EoE. Likewise, experimental EoE requires IL-13Rα1 and TSLP^16, 17^. Whether these two axes converge on shared epithelial programs, define distinct disease endotypes, or operate in parallel within the same tissue remains to be defined. A precedent for the latter possibility exists in experimental eosinophilic gastritis. In that model, IL-4Rα blockade abrogated both inflammatory cell infiltration and epithelial remodeling, whereas deletion of *Il13ra1* selectively suppressed remodeling without affecting infiltration^53^. Distinct receptor-driven pathways can therefore control separate components of a single tissue phenotype.

In summary, we define the cellular and transcriptional landscape of established experimental EoE. Th2 cells and a cycling mucosal mast cell state dominate type 2 cytokine expression, whereas an interferon-associated program operates in parallel both in immune and structural compartments, and is conserved in human disease. These data provide a cellular framework for interpreting cytokine-directed interventions in EoE and identify the mucosal mast cell transition and the resident memory-like CD8⁺ T cell compartment as the populations whose functional contribution remains to be defined.

## Materials and Methods

### Mice

Male wild-type (WT) C57BL/6 mice were purchased from Harlan Laboratories (Israel). *Il13ra1*^⁻/⁻^ mice on a C57BL/6 background were generated and bred in house^16^. All mice were housed under specific pathogen-free conditions at Tel Aviv University and were 6–8 weeks old at the start of experiments. All experiments were approved by the Tel Aviv University Institutional Animal Care and Use Committee.

### Experimental eosinophilic esophagitis

Experimental EoE was induced as previously described^14^. Mice were skin-sensitized with 1% oxazolone (OXA; Sigma, #E0753) in acetone applied to both ears (60 µL per mouse), followed by five skin challenges with 0.5% OXA in acetone. From day 18, mice received a total of eight intraesophageal challenges with 200 µL of 1% OXA in a 1:2 mixture of olive oil and 95% ethanol, delivered through a plastic feeding tube reaching the mid-esophagus (INSTECH, #ITH-FTP-22-25-50, 22 ga × 25 mm) that had been modified for intraesophageal administration by puncturing eight holes with a 14G needle. Control mice were left untreated (naive; single-cell experiment) or received vehicle only (histology experiment). Esophagi were collected on day 36.

### Histology, immunohistochemistry and immunofluorescence

Esophagi were fixed in 4% buffered formaldehyde, processed on a Leica tissue processor, embedded in paraffin and cut into 5-µm sections. Sections were stained with hematoxylin and eosin, and epithelial thickness was measured with the line tool in QuPath (six measurements per section, four sections per mouse, averaged per mouse). Eosinophils were detected by immunohistochemistry for major basic protein (anti-MBP; antibody kindly provided by Dr. Elizabeth A. Jacobsen, Mayo Clinic) and counted per section. Proliferating cells were detected with a rabbit anti-Ki67 antibody (Novus Biologicals, #NB500-170; 1:750; heat-induced epitope retrieval with BOND Epitope Retrieval Solution 1) on a Leica BOND automated Stainer and expressed per mm² of epithelium. Mucosal mast cells were detected with a rabbit monoclonal anti-mast cell protease 1 antibody (MCPT1; clone EPR25920A-86, Abcam; 1:500) on a Leica BOND automated stainer. Keratin 6 (KRT6; BioLegend, #905701; 1:1000) and keratin 14 (KRT14; BioLegend, #905301; 1:1000) were detected by immunofluorescence using conjugated secondary antibodies with DAPI nuclear counterstaining. Slides were scanned on a Leica Aperio slide scanner. Scale bars represent 50 µm. Quantification was performed blinded to group on four sections per mouse.

### Preparation of single-cell suspensions and single-cell RNA sequencing

Esophagi from naive and OXA-challenged (day 36) WT mice were pooled (2-4 esophagi per sample; two pooled samples per condition). Tissue was minced and digested with collagenase type IV (Sigma-Aldrich, #C5138 and DNase I (Roche, #10104159001) for 40 min at 37 °C with shaking at 200 rpm, and filtered through a 40-µm strainer. Dead cells were removed with the Dead Cell Removal Kit (Miltenyi Biotec, #130-090-101), and CD45⁺ cells were enriched with mouse CD45 MicroBeads (Miltenyi Biotec, #130-052-301) on LS columns (Miltenyi Biotec, #130-042-401). This enrichment retained a fraction of CD45⁻ structural cells. For each sample, approximately 60,000 cells were loaded on a separate BD Rhapsody cartridge, and whole-transcriptome libraries were prepared with the BD Rhapsody WTA Amplification Kit (BD Biosciences) according to the manufacturer’s instructions. Libraries were sequenced on an Illumina NovaSeq X (paired-end; read 1, 51 bp; read 2, 71 bp).

### Read processing and quality control

Reads were processed with the BD Rhapsody Sequence Analysis Pipeline on the Seven Bridges Genomics platform against the mouse GRCm39 reference genome. Count matrices were analyzed in R (v4.2.2) with Seurat (v5.0.3)^57^. Cells with more than 200 detected genes, more than 500 unique molecular identifiers (UMIs) and less than 20% mitochondrial reads were retained; the permissive mitochondrial threshold reflects the high mitochondrial content of esophageal cells. These criteria retained 71,268 of 77,222 cells (92.3%). Doublets were identified for each sample with scDblFinder (v1.12.0)^58^ on raw counts with default settings and removed (9,023 cells, 12.7%; 9.5-16.3% per sample), leaving 62,245 cells (33,544 from naive and 28,701 from OXA-challenged samples).

### Normalization, integration, clustering and annotation

Counts were log-normalized (scale factor 10,000), 2,000 variable genes were selected (variance-stabilizing transformation)^59^, data were scaled and 30 principal components were calculated. Samples were integrated with Harmony (v1.2.0)^60^ on sample identity, and UMAP embeddings and shared nearest-neighbour graphs were computed from 30 Harmony dimensions. Clusters (resolution 0.6) were annotated using canonical marker genes and cluster markers (Wilcoxon rank-sum test, positive markers) and assigned to an immune (34,946 cells) or a structural (26,928 cells) compartment; 371 cells with mixed profiles were excluded. Each compartment was re-embedded separately. Lymphoid cells were re-embedded with 3,000 variable genes, 40 principal components and Harmony (UMAP of 30 dimensions, minimum distance 0.35) and annotated into 13 populations. Myeloid cells (18,534 cells) were re-embedded with 3,000 variable genes excluding ribosomal, mitochondrial, sex-linked, hemoglobin, Gm and stress-response genes, 40 principal components and Harmony, and clustered without prior labels at resolution 0.5; two mucosal mast cell clusters were merged, 27 basophils were set aside, and the resulting 16 clusters were annotated by marker genes. A spinous epithelial cluster (1,169 cells) was not shown in the figures.

### Cell composition, expression plots and statistics of single-cell data

Population frequencies were calculated for each pooled sample within the lymphoid or myeloid compartment. Frequencies of structural populations were not analyzed, because the fraction of structural cells depended on the efficiency of CD45⁺ enrichment. Bars show the mean of the two samples per condition and error bars the SEM. Differences were tested with two-tailed unpaired *Student’s t tests* without correction for multiple comparisons and are considered descriptive given two pooled samples per condition. In dot plots, dot size indicates the percentage of cells with at least one detected transcript, and colour indicates either the average expression scaled across groups (z-score) or the mean expression relative to the highest group, as stated in the figure legends. Heat maps of individual samples show row z-scores of pseudobulk log counts per million (CPM). Cell-level tests pooled the cells of both samples per condition; because cells from the same sample are not independent, these P values were used to rank genes and not to infer differences between animals.

### Type 2 program score

A type 2 effector and repair program (*Il13*, *Il4*, *Il5*, *Areg*, *Gata3* and *Il1rl1*) was scored in lymphoid cells with ModuleScore (Seurat; seed 42)^61^. Program-high cells were defined as cells with a score above the 90th percentile of all lymphoid cells in which at least one of *Il4*, *Il5* or *Il13* was detected. The same genes were scored in lymphoid cells of a published human EoE atlas (Broad Single Cell Portal SCP1242^12^; healthy individuals, patients in remission and patients with active disease), using the authors’ annotations and the mean of per-gene z-scores, with the same two criteria. For the specificity comparison (Fig. S1C), each gene was given the percentile rank (0 to 1) of its mean expression in Th2 cells among all immune cell types of the respective dataset, and program scores are the mean over the genes of each program.

### Differential expression

Differential expression was calculated with the Wilcoxon rank-sum test (FindMarkers, Seurat) on log-normalized data with Bonferroni-adjusted P values. Th2 cells were compared with all other lymphoid cells and with effector CD4 T cells at day 36 (genes detected in at least 5% of cells in either group; Fig. 3A). Eosinophils were compared with all other myeloid cells, including neutrophils, at day 36 (Fig. S4; dashed lines, |log2 fold change| = 1 and adjusted P = 0.05). Mucosal-type mast cells were compared with connective-type mast cells at day 36.

### Gene set enrichment analysis

Ranked gene set enrichment analysis^62^ was performed with fgsea^63^ (multilevel, 10,000 simple permutations) using Hallmark^64^, Reactome^65^, KEGG^66^ and Gene Ontology^67^ biological process gene sets (MSigDB, mouse orthologues via msigdbr; size limits 15-300 genes for Gene Ontology and 10-400 genes otherwise), and P values were adjusted by the Benjamini-Hochberg method. Genes were ranked by a rank-based statistic of Th2 cells versus all other lymphoid cells at day 36 (Fig. 3B); by a pseudobulk statistic of EoE versus naive samples for ILC2s, regulatory T cells and CD8⁺ T cells (Fig. S2); by mucosal-type versus connective-type mast cells (Fig. 6C); and, for epithelial cells (Fig. 7C), by the signed -log10 P value of the Wilcoxon test of day 36 versus naive cells in each layer, excluding upregulated genes whose expression could be explained by ambient RNA from immune cells (mean expression at day 36 not exceeding three times the expected ambient contribution).

### Macrophage program scores

For inflammatory (C1), Lyve1⁺Folr2⁺ (C2) and C1q⁺Cx3cr1⁺ (C3) macrophages, pseudobulk log2 CPM was calculated for each sample. Genes of ten predefined functional programs detected in at least 5% of cells were scaled from 0 to 1 across the 12 population-by-sample values, and the program score was represented as the mean over the program genes.

### Mast cell classification

Mast cells were classified by protease expression^68^. Cells were classified as mucosal-type if the mean log-normalized expression of *Mcpt1*, *Mcpt2* and *Mcpt9* was at least 0.5 or both *Mcpt1* and *Mcpt2* were detected, and as connective-type otherwise. Cells with detected *Mki67* or *Top2a* were classified as cycling.

### Ligand-receptor analysis

Candidate interactions were derived from the mouse ligand-receptor network of NicheNet. Interaction strength was calculated as the product of the mean (non-log) normalized expression of the ligand in the sender population and of the receptor in the receiving population, shown as log10(1 + product). For Th2 cells, ligands were required to be detected in at least 10% of Th2 cells and enriched relative to other lymphoid cells, and receptors to be detected in at least 10% of receiving cells. For eosinophils, ligands from neighboring populations were paired with receptors detected in at least 10% of eosinophils in both samples. For mast cells, interactions were retained when both the mast cell ligand and the receptor on the receiving population increased in EoE (Wilcoxon test, adjusted P < 0.05, at least 1.2-fold), the receptor was detected in at least 10% of receiving cells and the receiving population contained at least 30 cells in naive samples. For non hematopoietic cells, ligands that increased in epithelial cells or fibroblasts in EoE were paired with receptors on immune populations. These are predictions based on co-expression and were not validated functionally.

### Comparison with published datasets

Eosinophil-enriched genes were compared by Spearman correlation of log2 fold changes with an independent mouse esophagus dataset (GSE262177^22^; eosinophils versus all other cells) and with esophageal eosinophils from six patients with active EoE (GSE175930^23^; the eosinophil cluster versus other myeloid cells, recomputed in this study). Human genes were mapped to mouse orthologues with HCOP^23^ (babelgene). Overlaps with genes induced in eosinophils in other settings (Fig. 5G) were expressed as fold enrichment over the overlap expected by chance (hypergeometric test). These datasets were lung metastasis (GSE181491)^17^, colitis-associated colon tumours (GSE112922)^27^, *Citrobacter rodentium* colitis and *Helicobacter pylori* gastritis (GSE182001)^28^, atopic dermatitis skin (GSE224703)^24^, *in vitro* type 1 and type 2 polarization (GSE243369)^29^, IL-33-induced bone marrow eosinophilia (GSE229824)^69^ and allergic lung TREM-1⁺ eosinophils (GSE246389)^26^. Human esophageal biopsies of GSE201153^30^ were used. Epithelial pathways were analyzed in five patients with active EoE and two healthy controls (pseudobulk per patient for basal and differentiated epithelial cells, genes ranked by log2 fold change active versus healthy). Pathways were classified as shared when significant in the same direction in at least one layer of each species and never opposite, and as specific to one species when significant there and either not significant (FDR > 0.25) or opposite in both layers of the other species. Human mast cells (active EoE biopsies) and mouse mast cells (day 36) were each compared with an equal-weight reference of T/NK cells and myeloid antigen-presenting cells, and log2 fold changes were compared for 3,622 one-to-one orthologues.

### Statistical analysis

Histology data are shown as mean ± SEM with each dot representing one mouse (n = 8-10 mice per group) and were analyzed by two-way ANOVA with Tukey’s multiple comparisons test in GraphPad Prism (v10). Single-cell analyses are based on two pooled samples (2-4 mice each) per condition. P < 0.05 was considered significant.

## Software and data availability

Analyses used R (v4.2.2) with Seurat (v5.0.3), harmony (v1.2.0), scDblFinder (v1.12.0), fgsea, msigdbr, babelgene, edgeR^70^ and ggplot2 (v3.5.1). The single-cell data have been deposited in the Gene Expression Omnibus.

