## Supplemental Data for "Single-cell analysis identifies concurrent type 2 and interferon programs in experimental eosinophilic esophagitis"

**Short Title:** Single cell analysis of experimental EoE

***Corresponding authors:** Ariel Munitz, PhD, The Roberts-Guthman Chair in Immunopharmacology, Department of Clinical Microbiology and Immunology, Gray Faculty of Medical and Health Sciences, Tel-Aviv University, Israel. Ramat Aviv 69978, Israel. Tel. (Office): +972-3-640-7636, Fax: +972-3-640-9160,; ORCID: https://orcid.org/0000-0003-1626-3019; Anish Dsilva,

MSc, Department of Clinical Microbiology and Immunology, Gray Faculty of Medical and Health Sciences, Tel-Aviv University, Israel. Ramat Aviv 69978, Israel,; ORCID: https://orcid.org/0009-0002-4898-6086

**Grant Support:** Ariel Munitz is supported by the US-Israel Bi-national Science Foundation (grant no. 2023029), the Food Allergy Fund, the Israel Science Foundation (grants no. 542/20) The Israel Science Foundation-Swiss National Science Foundation Collaborative Grant 954/26 (954/26 with ICA), the Israel Cancer Research Fund, the Israel Cancer Association Avraham Rotstein Donation, the Cancer Biology Research Center (TAU), the Azrieli Foundation Canada-Israel and External sponsored scientific research grants from AstraZeneca.

**Disclosures:** Ariel Munitz is a consultant for Glaxo Smith Kline, Astra Zeneca, Sanofi and is an inventor of patents owned by the Tel Aviv University.

### Figure legends

**Figure S1. Th2 cells predominate among type 2 program-high lymphocytes in human and mouse EoE.**

Lymphoid cells of the human EoE single-cell atlas (SCP1242) were analyzed using the cell annotations of the single cell portal (https://singlecell.broadinstitute.org/single_cell), and type 2 program-high cells from control individuals, patients in remission and patients with active EoE are shown (A). Type 2 program-high cells of naive and EoE mice are shown on the mouse lymphoid UMAP (B). Program-high cells were defined by a mean z-score comprised of of *IL13*, *IL4*, *IL5*, *AREG*, *GATA3* and *IL1RL1* is shown. The specificity of Th2 programs in human and mouse is presented (C). For each gene, the percentile rank (0 to 1) of mean expression in Th2 cells among all immune cell types was determined and averaged over the genes of each program.

**Figure S2. Interferon and inflammatory pathways are induced in ILC2s, regulatory T cells and $CD8^+$ T cells in experimental EoE.**

GSEA of ILC2s (A), $CD4^+$ regulatory T cells (C) and $CD8^+$ T cells (E) of EoE versus naive mice is shown. Pale bars are nonsignificant. Expression of leading-edge genes of selected pathways in each sample is presented as row z-scores of pseudobulk log counts per million (B, D, F); 2*adjusted* $p < 0.05$, *adjusted* $p < 0.01$, *adjusted* $p < 0.001$.

**Figure S3. Differentialy expressed genes in macrophage populations.**

Expression of the genes that define each functional program of inflammatory (S3-left panel), Lyve1/Folr2 (S3-middle panel) and C1q/Cx3cr1 (S30right panel) macrophages is shown for each naive and EoE sample as row z-scores of pseudobulk log2 counts per million. Grey color indicates a gene that was not detected in that population.

**Figure S4. Eosinophil-enriched genes in experimental EoE.**

Genes differentially expressed between eosinophils and all other myeloid cells of EoE mice (Wilcoxon test) are shown as a volcano plot. Dashed lines represent a log2 fold change of ±1 and an adjusted p value of 0.05. Selected genes are labeled and colored by category.

**Figure S5. Mast cell states, signals and comparison with human EoE.**

Genes differentially expressed between mucosal-type and connective-type mast cells of EoE mice are shown as a volcano plot (A; x-axis compressed above a log2 fold change of 4). The percentage of sender cells expressing ligands that can reach mast cells in naive and EoE mice, and the percentage of connective-type/mucosal-type mast cells expressing the receptor, are presented (B). Open circles represent individual samples and filled circles the mean. Ligands expressed by mast cells (percentage of mast cells and fold enrichment over other immune cells) and the expression of their receptors in receiving cells are shown (C). Dot size represents the percentage of receiving cells expressing the receptor and dot color the interaction strength. Mast cell-specific genes in human and mouse EoE are compared (D). Each dot represents a gene with a one-to-one mouse orthologue, and axes show the log2 fold change of mast cells versus an equal-weight reference of T/NK cells and myeloid antigen-presenting cells in active human EoE biopsies (GSE201153; x-axis) and in experimental EoE (y-axis); rho, Spearman correlation.

Figure S1.

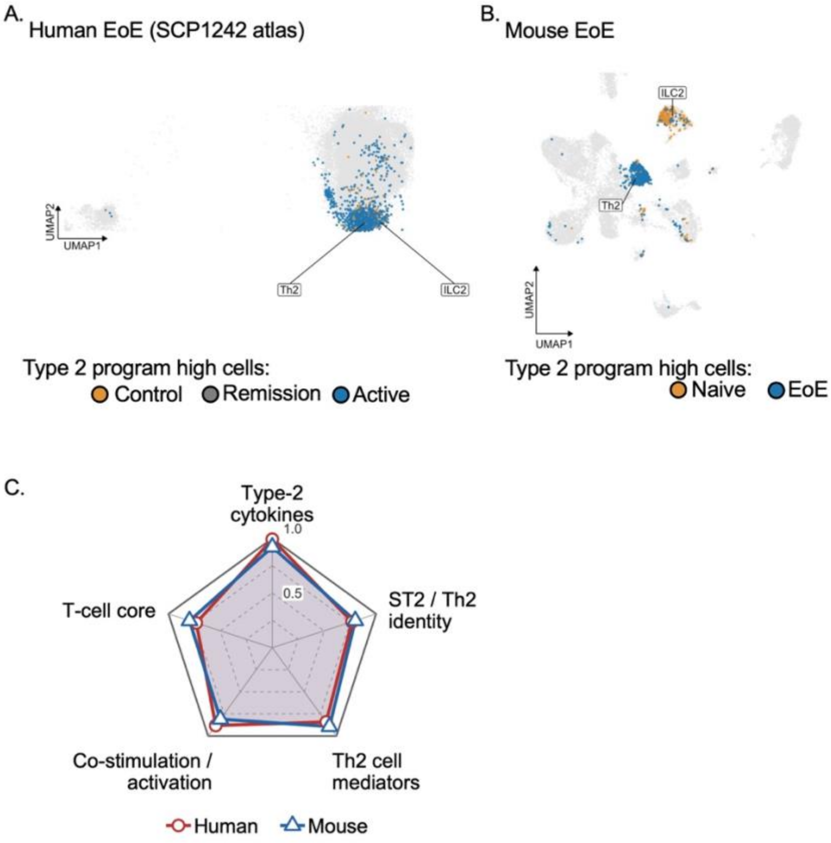
A. Human EoE (SCP1242 atlas)
UMAP2
UMAP1
Th2
ILC2
Type 2 program high cells:
Control
Remission
Active
B. Mouse EoE
ILC2
Th2
UMAP2
UMAP1
Type 2 program high cells:
Naive
EoE
C.
Type-2 cytokines
1.0
0.5
ST2 / Th2 identity
T-cell core
Co-stimulation / activation
Th2 cell mediators
Human
Mouse

Figure S2.

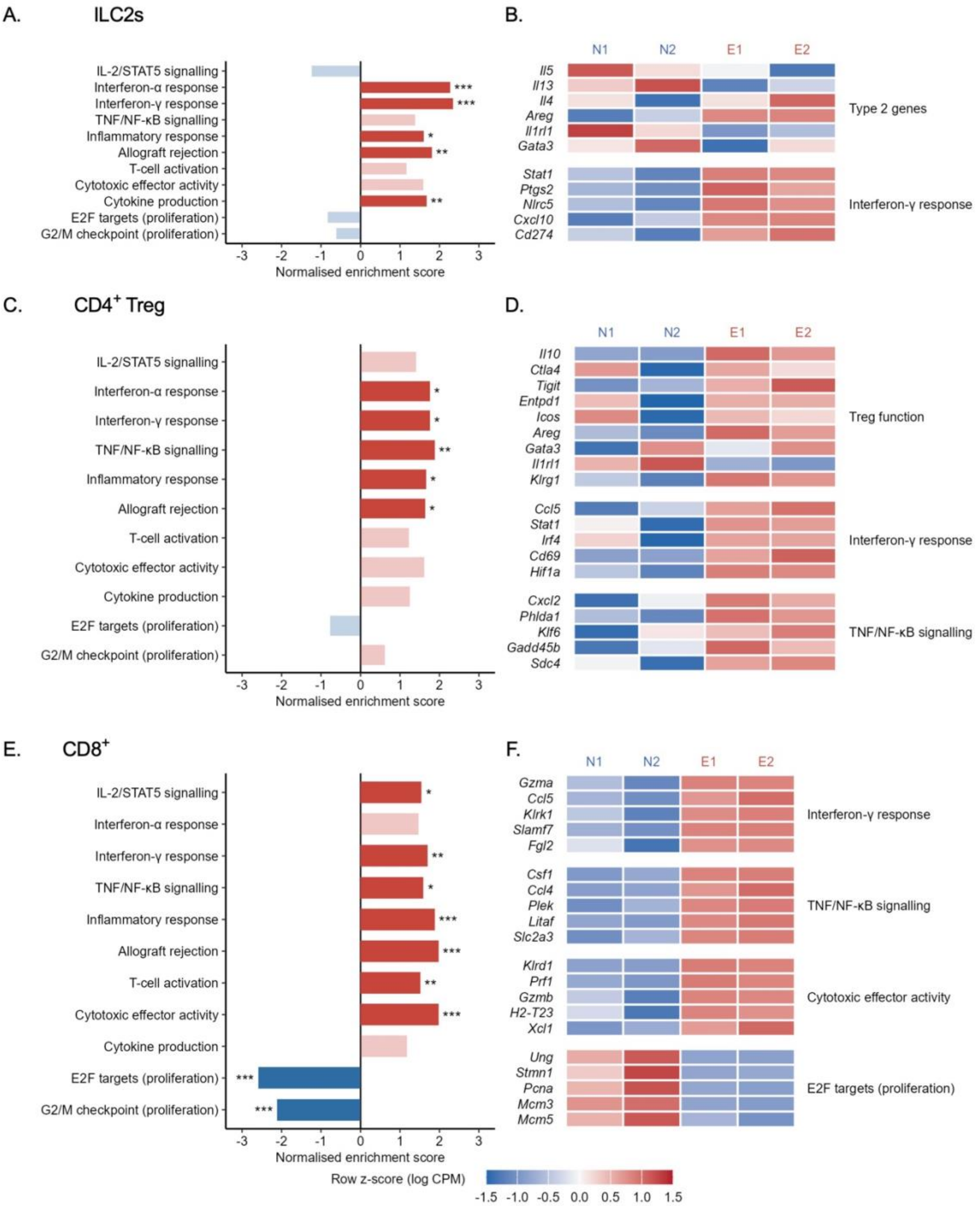

A.
ILC2s
IL-2/STAT5 signalling
Interferon-α response
Interferon-γ response
TNF/NF-κB signalling
Inflammatory response
Allograft rejection
T-cell activation
Cytotoxic effector activity
Cytokine production
E2F targets (proliferation)
G2/M checkpoint (proliferation)
Normalised enrichment score
B.
N1
N2
E1
E2
Il5
Il13
Il4
Areg
Il1rl1
Gata3
Type 2 genes
Stat1
Ptgs2
Nlrc5
Cxcl10
Cd274
Interferon-γ response
C.
CD4+ Treg
D.
Il10
Ctla4
Tigit
Entpd1
Icos
Areg
Gata3
Il1rl1
Klrg1
Treg function
Ccl5
Stat1
Irf4
Cd69
Hif1a
Cxcl2
Phlda1
Klf6
Gadd45b
Sdc4
TNF/NF-κB signalling
E.
CD8+
F.
Gzma
Ccl5
Klrk1
Slamf7
Fgl2
Csf1
Ccl4
Plek
Litaf
Slc2a3
Klrd1
Prf1
Gzmb
H2-T23
Xcl1
Cytotoxic effector activity
Ung
Stmn1
Pcna
Mcm3
Mcm5
E2F targets (proliferation)
Row z-score (log CPM)
-1.5 -1.0 -0.5 0.0 0.5 1.0 1.5

Figure S3.

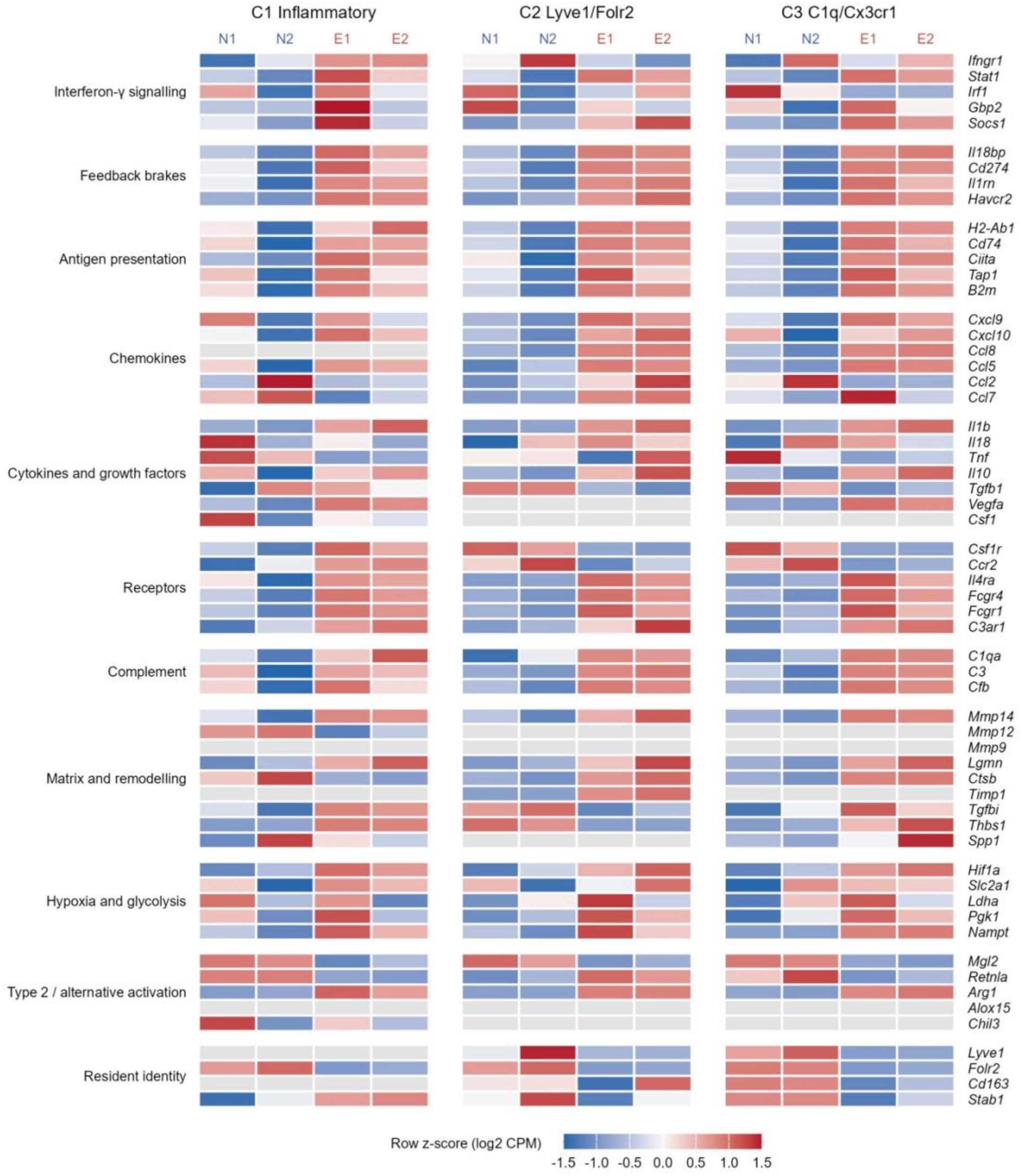
C1 Inflammatory
C2 Lyve1/Folr2
C3 C1q/Cx3cr1
N1
N2
E1
E2
Interferon-γ signalling
Feedback brakes
Antigen presentation
Chemokines
Cytokines and growth factors
Receptors
Complement
Matrix and remodelling
Hypoxia and glycolysis
Type 2 / alternative activation
Resident identity
Ifngr1
Stat1
Irf1
Gbp2
Socs1
Il18bp
Cd274
Il1rn
Havcr2
H2-Ab1
Cd74
Ciita
Tap1
B2m
Cxcl9
Cxcl10
Ccl8
Ccl5
Ccl2
Ccl7
Il1b
Il18
Tnf
Il10
Tgfb1
Vegfa
Csf1
Csf1r
Ccr2
Il4ra
Fcgr4
Fcgr1
C3ar1
C1qa
C3
Cfb
Mmp14
Mmp12
Mmp9
Lgmn
Ctsb
Timp1
Tgfbi
Thbs1
Spp1
Hif1a
Slc2a1
Ldha
Pgk1
Nampt
Mgl2
Retnla
Arg1
Alox15
Chil3
Lyve1
Folr2
Cd163
Stab1
Row z-score (log2 CPM)
-1.5 -1.0 -0.5 0.0 0.5 1.0 1.5

Figure S4.

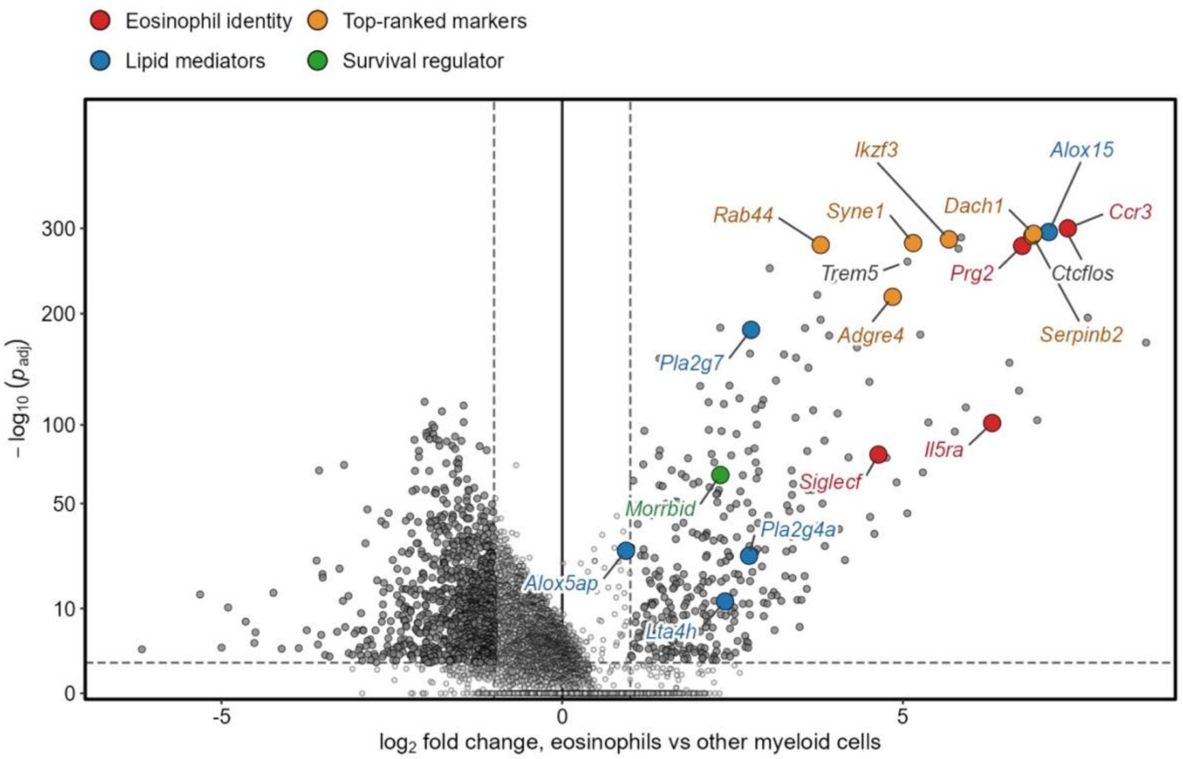

Eosinophil identity
Top-ranked markers
Lipid mediators
Survival regulator
Ikzf3
Alox15
Rab44
Syne1
Dach1
Ccr3
Trem5
Prg2
Ctcflos
Adgre4
Serpinb2
Pla2g7
Il5ra
Siglecf
Morrbid
Pla2g4a
Alox5ap
Lta4h
300
200
100
50
10
0
-5
0
5
$-\log_{10}(p_{adj})$
$\log_2$ fold change, eosinophils vs other myeloid cells

Figure S5.

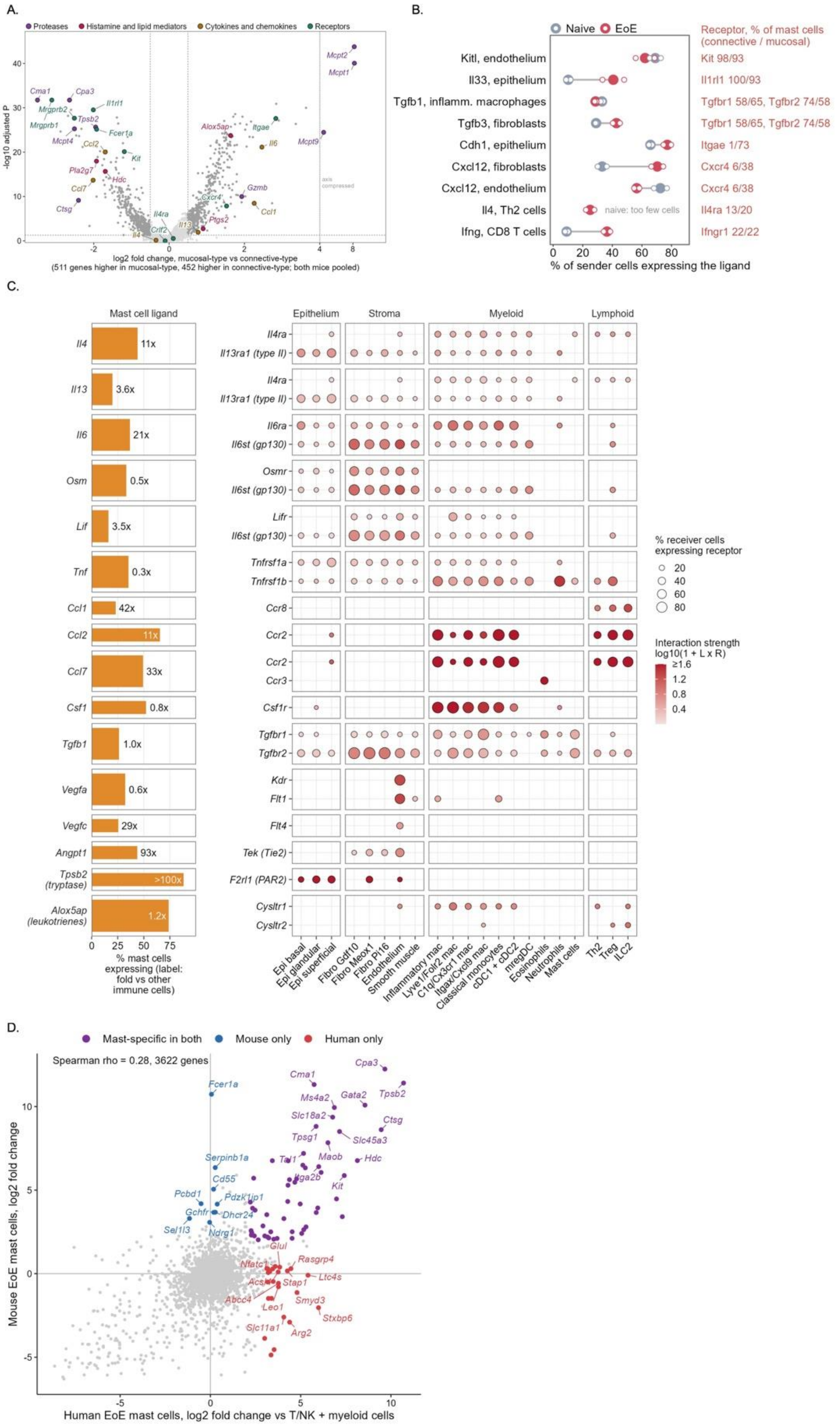

A.
Proteases
Histamine and lipid mediators
Cytokines and chemokines
Receptors
-log10 adjusted P
log2 fold change, mucosal-type vs connective-type
(511 genes higher in mucosal-type, 452 higher in connective-type; both mice pooled)
Mcpt2
Mcpt1
Mcpt9
Cma1
Cpa3
Mrgprb2
Mrgprb1
Il1rl1
Tpsb2
Fcer1a
Mcpt4
Ccl2
Kit
Pla2g7
Hdc
Ccl7
Ctsg
Alox5ap
Itgae
Il6
Gzmb
Cxcr4
Ccl1
Ptgs2
Il4ra
Il13
Crlf2
Il4
axis compressed
B.
Naive
EoE
Receptor, % of mast cells (connective / mucosal)
Kitl, endothelium
Il33, epithelium
Tgfb1, inflamm. macrophages
Tgfb3, fibroblasts
Cdh1, epithelium
Cxcl12, fibroblasts
Cxcl12, endothelium
Il4, Th2 cells
Ifng, CD8 T cells
Kit 98/93
Il1rl1 100/93
Tgfbr1 58/65, Tgfbr2 74/58
Tgfbr1 58/65, Tgfbr2 74/58
Itgae 1/73
Cxcr4 6/38
Cxcr4 6/38
Il4ra 13/20
Ifngr1 22/22
naive: too few cells
% of sender cells expressing the ligand
C.
Mast cell ligand
Epithelium
Stroma
Myeloid
Lymphoid
Il4 11x
Il13 3.6x
Il6 21x
Osm 0.5x
Lif 3.5x
Tnf 0.3x
Ccl1 42x
Ccl2 11x
Ccl7 33x
Csf1 0.8x
Tgfb1 1.0x
Vegfa 0.6x
Vegfc 29x
Angpt1 93x
Tpsb2 (tryptase) >100x
Alox5ap (leukotrienes) 1.2x
% mast cells expressing (label: fold vs other immune cells)
Il4ra
Il13ra1 (type II)
Il6ra
Il6st (gp130)
Osmr
Lifr
Tnfrsf1a
Tnfrsf1b
Ccr8
Ccr2
Ccr3
Csf1r
Tgfbr1
Tgfbr2
Kdr
Flt1
Flt4
Tek (Tie2)
F2rl1 (PAR2)
Cysltr1
Cysltr2
Epi basal
Epi glandular
Epi superficial
Fibro Gdf10
Fibro Meox1
Fibro Pi16
Endothelium
Smooth muscle
Inflammatory mac
Lyve1/Folr2 mac
C1q/Cx3cr1 mac
Itgax/Cxcl9 mac
Classical monocytes
cDC1 + cDC2
mregDC
Eosinophils
Neutrophils
Mast cells
Th2
Treg
ILC2
% receiver cells expressing receptor
Interaction strength log10(1 + L x R)
D.
Mast-specific in both
Mouse only
Human only
Spearman rho = 0.28, 3622 genes
Mouse EoE mast cells, log2 fold change
Human EoE mast cells, log2 fold change vs T/NK + myeloid cells
Cpa3
Cma1
Fcer1a
Ms4a2
Gata2
Tpsb2
Slc18a2
Ctsg
Tpsg1
Slc45a3
Serpinb1a
Tal1
Maob
Hdc
Cd55
Itga2b
Kit
Pcbd1
Pdzk1ip1
Gchfr
Dhcr24
Sel1l3
Ndrg1
Glul
Nfatc1
Rasgrp4
Ltc4s
Acsl1
Stap1
Abcc4
Smyd3
Leo1
Stxbp6
Slc11a1
Arg2
